# Common genetic variation influencing human white matter microstructure

**DOI:** 10.1101/2020.05.23.112409

**Authors:** Bingxin Zhao, Tengfei Li, Yue Yang, Xifeng Wang, Tianyou Luo, Yue Shan, Ziliang Zhu, Di Xiong, Mads E. Hauberg, Jaroslav Bendl, John F. Fullard, Panagiotis Roussos, Yun Li, Jason L. Stein, Hongtu Zhu

**Author notes:** Corresponding author: Hongtu Zhu, 3105C McGavran-Greenberg Hall, 135 Dauer Drive, Chapel Hill, NC 27599. List of Pediatric Imaging, Neurocognition and Genetics (PING) authors provided in the supplemental materials.

## Abstract

Brain regions communicate with each other via tracts of myelinated axons, commonly referred to as white matter. White matter microstructure can be measured in the living human brain using diffusion based magnetic resonance imaging (dMRI), and has been found to be altered in patients with neuropsychiatric disorders. Although under strong genetic control, few genetic variants influencing white matter microstructure have ever been identified. Here we identified common genetic variants influencing white matter microstructure using dMRI in 42,919 individuals (35,741 in the UK Biobank). The dMRIs were summarized into 215 white matter microstructure traits, including 105 measures from tract-specific functional principal component analysis. Genome-wide association analysis identified many novel white matter microstructure associated loci (*P* < 2.3 × 10^−10^). We identified shared genetic influences through genetic correlations between white matter tracts and 62 other complex traits, including stroke, neuropsychiatric disorders (e.g., ADHD, bipolar disorder, major depressive disorder, schizophrenia), cognition, neuroticism, chronotype, as well as non-brain traits. Common variants associated with white matter microstructure alter the function of regulatory elements in glial cells, particularly oligodendrocytes. White matter associated genes were enriched in pathways involved in brain disease pathogenesis, neurodevelopment process, and repair of white matter damage (*P* < 1.5 × 10^−8^). In summary, this large-scale tract-specific study provides a big step forward in understanding the genetic architecture of white matter and its genetic links to a wide spectrum of clinical outcomes.

---

Brain functions depend on effective communication across brain regions^1^. White matter comprises roughly half of the human brain and contains most of the brain’s long-range communication pathways^2^. White matter tracts build a complex network of structural connections, which keeps the brain globally connected and shapes communication and connectivity patterns^3-5^. Cellular microstructure in white matter tracts plays a pivotal role in maintaining the integrity of connectivity and mediating signal transitions among distributed brain regions^6^. Evidence from neuroscience has further suggested that white matter microstructure may underpin brain function and dysfunction^1,7,8^, and connectivity differences or changes are relevant to a wide variety of neurological and psychiatric disorders, such as attention-deficit/hyperactivity disorder (ADHD)^9^, major depressive disorder (MDD)^10^, schizophrenia^11^, bipolar disorder^12^, multiple sclerosis^13^, Alzheimer’s disease^14^, corticobasal degeneration^15^, and Parkinson’s disease^16^. White matter microstructural differences and abnormalities can be captured *in vivo* by diffusion magnetic resonance imaging (dMRI). Using dMRI data, microstructural connectivity can be quantified in diffusion tensor imaging (DTI) models^17^ and measured by several DTI-derived parameters, including fractional anisotropy (FA), mean diffusivity (MD), axial diffusivity (AD), radial diffusivity (RD), and mode of anisotropy (MO). Among them, FA serves as the primary metric of interest in many studies^18^, which is a robust global measure of integrity/directionality and is highly sensitive to general connectivity changes. On the other hand, MD, AD, and RD directly quantify the abstract magnitude of directionalities, and thus are more sensitive to specific types of microstructural changes^19^. In addition, MO can characterize the anisotropy type, describing whether the shape of the diffusion tensor is more linear or planar^20,21^. See **Supplementary Note** for a global overview of these commonly used DTI parameters.

White matter differences in general population cohorts are under strong genetic control. Both family and population-based studies have reported that DTI measurements of white matter microstructure have in general high heritability with estimates varying across different age groups^22^ and tracts^23^. For example, heritability estimates of tract-averaged FA ranged from 53% to 90% in twin study of the Human Connectome Project (HCP)^24^. Recent genome-wide association studies (GWAS) of UK Biobank reported an average SNP-based heritability of 48.7% across different tracts^25^.

Several GWAS^23,25-29^ have been performed to identify loci associated with inter-individual variation in white matter microstructure but shared at least two major limitations: (i) sample size and (ii) spatial specificity. First, the current largest published GWAS of dMRI phenotypes has sample size 17,706 in Zhao, et al. ^25^. Similar to other brain-related traits^30^, white matter has a complex and extremely polygenic genetic architecture^25,31^. Large sample size is essential to boost GWAS power in order to identify many common risk variants with small effect sizes. Second, previous GWAS mainly focused on global dMRI measures of the whole brain^26,27^ or tract-averaged (mean) values^23,25^. Global and tract-averaged measures can capture the largest variations in white matter, while reducing the burden to test multiple neuroimaging traits, particularly suitable for GWAS with limited sample size; however, these measures may lose lots of information, as microstructural differences and changes may not have a uniformly consistent pattern across the whole tract. Heterogeneous variation patterns typically exist within voxel-wise DTI maps of the 3D tract curve, which may be more relevant to specific underlying biological processes. For example, previous study found that the association between bipolar disorder and FA is specific to one given segment of the long anterior limb of internal capsule (ALIC) tract connecting prefrontal cortex with the thalamus and brain stem^32^. Due to these limitations, a large number of genetic factors influencing white matter may still be undiscovered. Consequently, with few exceptions (e.g., stroke^26^ and cognitive traits^25^), the shared genetic influences between white matter and other complex traits are unknown. Uncovering these potential genetic links may identify important brain regions that are involved in clinical outcomes, especially for brain-related disorders.

To overcome these limitations, here we collected individual-level dMRI from five data resources: the UK Biobank^33^, Adolescent Brain Cognitive Development (ABCD^34^), HCP^35^, Pediatric Imaging, Neurocognition, and Genetics (PING^36^), and Philadelphia Neurodevelopmental Cohort (PNC^37^). We harmonized image processing by using the ENIGMA-DTI pipeline^38,39^ and obtained voxel-wise DTI maps for 42,919 subjects (after quality controls), including 35,741 in UK Biobank. We mainly focused on 21 predefined white matter tracts and generated two groups of phenotypes. The first group contains 110 tract-averaged parameters for FA, AD, MD, MO and RD in 21 tracts and across the whole brain. Second, we applied functional principal component analysis (FPCA^40^) to generate 105 tract-specific principal components (PCs) for FA by taking the top five PCs of the voxel-wise map within each tract. FPCA is a data-driven approach to characterize the strongest variation components of FA within each tract, which are expected to provide additional microstructural details about axonal organization and myelination omitted by tract-averaged values^41,42^, while limiting multiple testing. More importantly, these PCs may represent FA changes that are more relevant to specific clinical outcomes. We then performed a genome-wide association analysis for these 215 phenotypes to discover the genetic architecture of white matter and explore the genetic links to a plethora of clinical endpoints in different trait domains. Our GWAS results have been made publicly available at https://github.com/BIG-S2/GWAS and can be easily browsed through our Brain Imaging Genetics Knowledge Portal (BIG-KP) https://bigkp.web.unc.edu/.

## RESULTS

### GWAS Discovery and Validation for 215 DTI parameters

Our discovery analysis utilized data from UKB subjects of British ancestry (*n* = 33,292). All of the 110 DTI mean parameters had significant SNP heritability^43^ (*h*^*2*^) after Bonferroni adjustment (215 tests, *P* < 9.4 × 10^−31^, **Fig. 1a** and **Supplementary Table 1**). The *h*^*2*^ estimates varied from 24.8% to 65.4% (mean *h*^*2*^ = 46.3%), which were comparable with previous results^23,25^. For the 105 tract-specific FA PC parameters, we found that 102 had significant *h*^*2*^ (mean *h*^*2*^ = 34.1%, *h*^*2*^ range = (8.6%, 65.8%), *P* < 1.1 × 10^−5^). The 4^th^ PC of corticospinal tract (CST, 6.2%), 5^th^ PC of cingulum hippocampus (CGH, 4.4%), and 4^th^ PC of superior fronto-occipital fasciculus (SFO, 3.7%) had nominally significant *h*^*2*^ estimates (*P* < 0.03), which became insignificant after Bonferroni adjustment. The top five PCs in external capsule (EC) were highlighted in bottom panels of **Figure 1b**. Different from tract-averaged value, these PCs captured more specific FA variations in distinct subfields of EC, all of which had high *h*^*2*^ (mean *h*^*2*^ = 47.9%, *h*^*2*^ range = (42.9%, 52.6%), *P* < 1.8 × 10^−89^). Another illustration was given in **Supplementary Figure 1** for the PCs of superior longitudinal fasciculus (SLF). These *h*^*2*^ results show that the additional microstructural variations captured by unconventional tract-specific FA PCs are also generally under genetic control. As illustrated in later sections, those heritable local FA variation patterns may also have higher power to identify the shared genetic influences with other complex traits.

**Figure 1:**
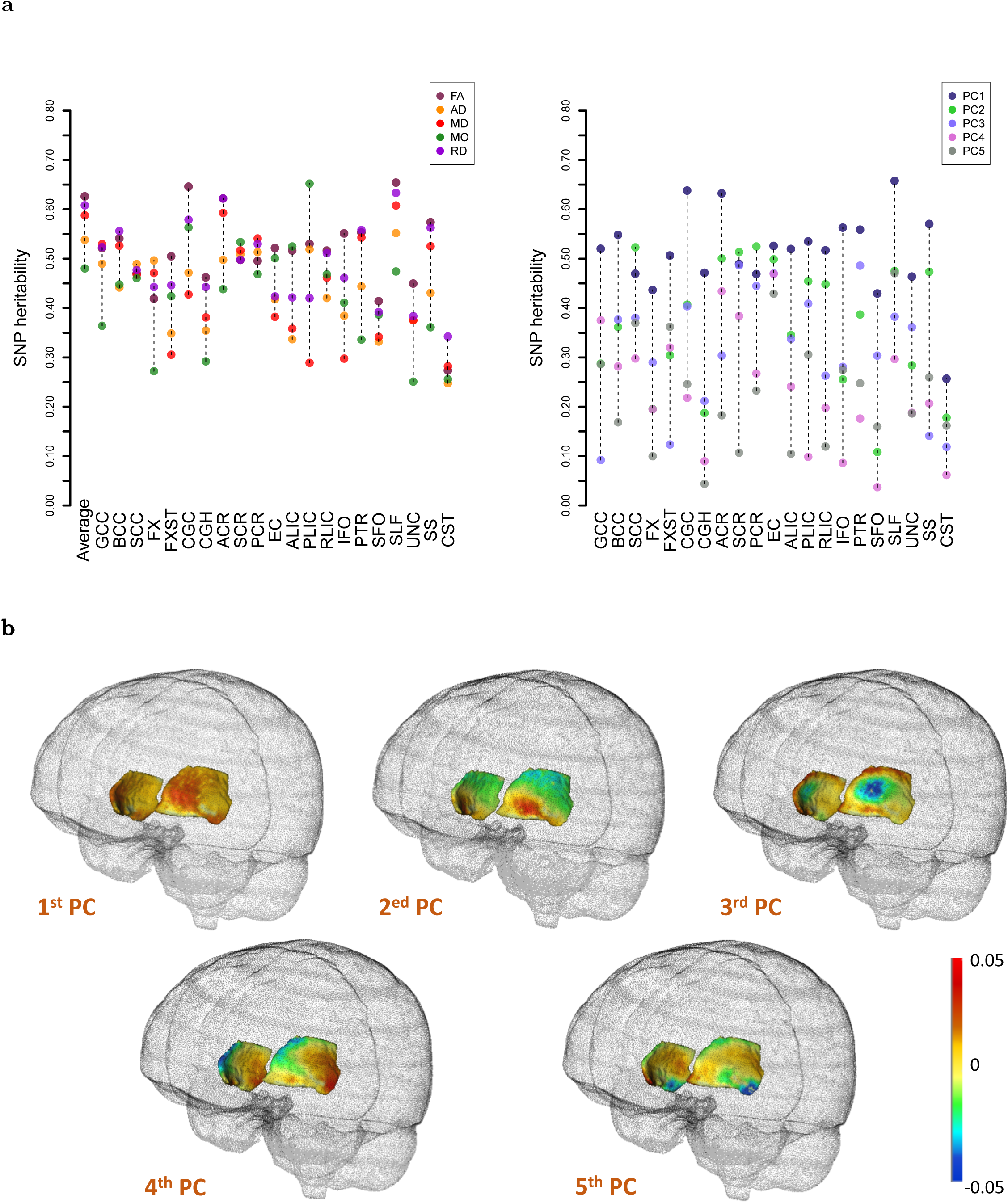
SNP heritability estimates of 215 DTI parameters (n = 33,292 subjects) and illustration of the top five FA principal components (PCs) of external capsule (EC). **a)** The 110 mean DTI parameters and 105 FA PC DTI parameters are displayed on the left and right panels, respectively. The x-axis lists the names of white matter tracts. **b)** The functional principal component (PC) loading coefficients for the top five FA PCs of EC.

We performed GWAS for these 215 DTI parameters using 9,023,710 common genetic variants after quality controls (Methods). All Manhattan and QQ plots can be browsed in our BIG-KP server. At a stringent significance level 2.3 × 10^−10^ (i.e., 5 × 10^−8^/215, additionally adjusted for the 215 phenotypes studied), FUMA^44^ clumped 595 partially independent significant variants (Methods) involved in 1,101 significant associations with 86 FA measures (21 mean and 65 PC parameters, **Supplementary Figs. 2-3** and **Supplementary Table 2**). Genetic variants had broad effects across all white matter tracts, and one variant often influenced multiple FA measures, such as rs12146713 in region 12q23.3, rs309587 in 5q14.3, rs55705857 in 8q24.21, and rs1004763 in 22q13.1. Of the 595 significant variants, 302 were only detected by PC parameters. On average, the number of FA-associated significant variants was 37.0 in each tract (range = (4, 72), **Fig. 2** and **Supplementary Table 3**), 50.3% of which were solely discovered by PC parameters (range = (26.3%, 100%)). For example, all of the 22 significant variants associated with CST were detected by PC parameters. Moreover, 66.7% (32/48) of the variants in posterior corona radiata (PCR), 64.9% (37/57) in posterior thalamic radiation (PTR), 59.7% (43/72) in SLF, and 56.3% (18/32) in cingulum cingulate gyrus (CGC) were only associated with PC parameters. These results clearly illustrate the unique contribution of tract-specific PC parameters in identifying genetic variants for FA variations within white matter tract.

**Figure 2:**
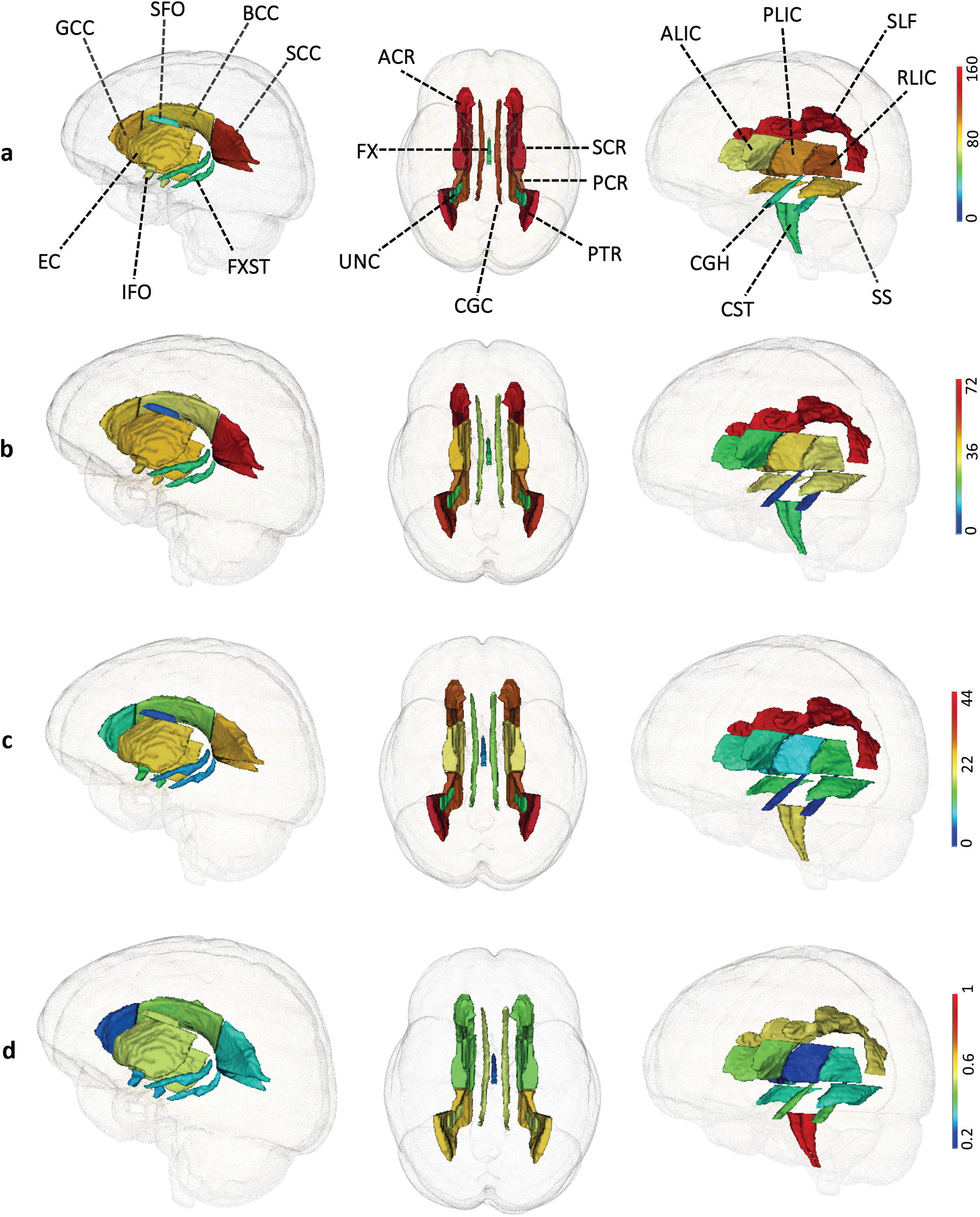
Number of independent significant variants identified in UKB British discovery GWAS at 2.3 × 10^−10^ significance level (n = 33,292 subjects). The first three rows are the number of independent significant variants identified in each white matter tract by **a)** any DTI parameters; **b)** any FA parameters; **c)** FA PC parameters, respectively. The last row **d)** displays the proportion of FA-associated variants that can only be identified by PC parameters.

In addition, 770 significant variants were associated with 83 mean parameters of AD, MD, MO and RD (2,069 significant associations), 565 of these 770 variants (with 967 associations) were not identified by FA measures (**Fig. 2, Supplementary Figs. 2-3**, and **Supplementary Table 2**). The mean number of significant variants in each tract moved up to 93.3 (range = (41, 160)), and rs13198474 in 6p22.2, rs2267161 in 22q12.2, rs55705857 in 8q24.21, rs7935166 in 11p11.2, and rs7225002 in 7q21.31 were associated with multiple non-FA measures. Of note, more than 70% of significant variants in cingulum (CGH (90.7%) and CGC (73.3%)) were detected by non-FA measures (**Supplementary Table 4**), which may suggest that FA is less useful in the thin line-like C-shaped cingulum region than in other tracts. Based on a second and more strict LD clumping (LD *r*^2^ < 0.1), FUMA^44^ defined independent lead variants from the above independent significant variants and then genetic loci were characterized (Methods). The 3,170 (1,101 + 2,609) significant variant-trait associations were summarized as 994 significant locus-trait associations (**Supplementary Tables 5-6**). We then performed functionally informed fine mapping for these locus-level signals using SuSiE^45^ via PolyFun^46^ framework (Methods). PolyFun + SuSiE identified 6,882 variant-trait pairs that had posterior causal probability (i.e., PIP) > 0.95 for 2,299 variants (**Supplementary Table 7**), suggesting the existence of multiple causal effects in associated loci. In summary, our results illuminate the broad genetics control on white matter microstructural differences. The genetic effects are spread across a large number of variants, consistent with the observed extremely polygenic genetic architecture of many brain-related traits^30,47^.

We aimed to find independent replication of our discovery GWAS in five independent validation datasets, all consisting of individuals of European ancestry: the UKB White but Non-British (UKBW, *n* = 1,809), ABCD European (ABCDE, *n* = 3,821), HCP (*n* = 334), PING (*n* = 461), and PNC (*n* = 537). First, for each DTI parameter, we checked the genetic correlation (gc) between discovery GWAS and the meta-analyzed European validation GWAS (total *n* = 6,962) by LDSC^48^ (Methods). The mean gc estimate was 0.95 (standard error = 0.35) across the 215 DTI parameters, 121 of which were significant after adjusting for multiple testing by the Benjamini-Hochberg (B-H) procedure at 0.05 level (**Supplementary Table 8**). Genetic correlation estimates near 1 indicates a consistent genetic basis for these phenotypes measured in different cohorts and MRI scanners. Next, we meta-analyzed our discovery GWAS with these European validation GWAS and found that 79.6% significant associations had smaller *P*-values after meta-analysis, suggesting similar effect size and direction of the top variants in independent cohorts^49,50^. Additionally, we tested for replication by using polygenic risk scores^51^ (PRS) derived from discovery GWAS (Methods). After B-H adjustment at 0.05 level (215 × 5 tests), the mean number of significant PRS in the five validation GWAS datasets was 195 (range = (193, 211), *P* range = (8.5 × 10^−27^, 4.5 × 10^−2^), **Supplementary Figs. 4-5** and **Supplementary Table 9**). Almost all (214/215) DTI parameters had significant PRS in at least one dataset and 165 had significant PRS in all of them, showing the high generalizability of our discovery GWAS results. Across the five validation datasets, the mean additional variance that can be explained by PRS (i.e., incremental R-squared) was 1.7% (range = (0.4%, 4.2%)) for the 165 consistently significant DTI parameters. The largest mean (incremental) R-squared was on the 2^nd^ PC of EC (range = (2.2%, 6.5%), *P* range = (7.2 × 10^−24^, 1.5 × 10^−9^)).

Finally, we constructed PRS on four non-European validation datasets: the UKB Asian (UKBA, *n* = 419), UKB Black (UKBBL, *n* = 211), ABCD Hispanic (ABCDH, *n* = 768), and ABCD African American (ABCDA, *n* = 1,257). The number of significant PRS was 158 and 40 in UKBA and UKBBL, respectively (B-H adjustment at 0.05 level, **Supplementary Table 10**). In addition, UKBW and UKBA had similar prediction performance (mean 2.38% vs. 2.33%, *P* = 0.67), but the accuracy became significantly smaller in UKBBL (mean 2.38% vs. 1.67%, *P* = 3.9 × 10^−9^). For the two non-European non-UKB datasets, the number of significant PRS was 121 and 114 in ABCDH and ABCDA, respectively (B-H adjustment at 0.05 level, **Supplementary Table 11**), which were much smaller than the ones observed in ABCDE. The R-squared were similar between ABCDH and ABCDE (mean 0.74% vs. 0.69%, *P* = 0.28), but the accuracy significantly decreased in ABCDA (mean 0.48% vs. 0.69%, *P* = 1.9 × 10^−7^). These findings show that UKB British GWAS findings have high generalizability in European cohorts, but the generalizability is reduced in cross-population applications, especially in Black/African-American cohorts, highlighting the importance of recruiting sufficient samples from global diverse populations in future genetics discovery of white matter.

### Concordance with previous GWAS

Of the 33,292 subjects in our UKB British discovery GWAS, 17,706 had been used in the largest previous GWAS^25^ for 110 mean parameters. To examine the robustness of their findings, we used the other 15,214 individuals (also removed the relatives^52^ of previous GWAS subjects) to perform a new validation GWAS and then evaluated the strength of replication (Methods). We calculated the replication slope, which was the correlation of the standardized effect size of variants estimated from two independent GWAS^53^. This analysis was restricted to top (*P* < 1 × 10^−6^ in previous GWAS) independent lead variants after LD-based clumping (window size 250, LD *r*^2^ = 0.01). The replication slope was 0.84 (standard error = 0.02, *P* < 2 × 10^−16^), indicating strong similarity between these top variant effect size estimates. We also applied FINDOR^53^ to reweight *P-*values by leveraging functional enrichments, after which the replication slope increased to 0.86 (standard error = 0.02, *P* < 2 × 10^−16^). In addition, for each of the 110 mean parameters, we used LDSC^48^ to calculate genetic correlation between measurements from the two GWAS. The mean gc estimate was 1.03 (standard error = 0.14, **Supplementary Fig. 6** and **Supplementary Table 12**) across these parameters, all of which were significant after B-H adjustment at 0.05 level (*P* < 1.4 × 10^−5^). In conclusion, these findings indicate that previous UKB GWAS results can be strongly validated in the new UKB British cohort.

Next, we carried out association lookups for 1,160 (595 + 565) independent significant variants (and variants within LD) detected in our UKB British discovery GWAS (Methods). Of the 213 variants (with 696 associations) identified in Zhao, et al. ^25^, 202 (with 671 associations) were in LD (*r*^2^ ≥ 0.6) with our independent significant variants (**Supplementary Table 13**). On the NHGRI-EBI GWAS catalog^54^, our results tagged many variants that had been implicated with brain structures, including 7 in van der Meer, et al. ^55^ for hippocampal subfield volumes, 7 in Verhaaren, et al. ^56^ for cerebral white matter hyperintensity (WMH) burden, 5 in Vojinovic, et al. ^57^ for lateral ventricular volume, 5 in Rutten-Jacobs, et al. ^26^ for WMH and white matter integrity, 2 in Klein, et al. ^58^ for intracranial volume, 2 in Hibar, et al. ^59^ for subcortical brain region volumes, 2 in Fornage, et al. ^28^ for WMH burden, 1 in Elliott, et al. ^23^ for brain imaging measurements, 1 in Luo, et al. ^60^ for voxel-wise brain imaging measurement, 1 in Hashimoto, et al. ^61^ for superior frontal gyrus grey matter volume, 1 in Ikram, et al. ^62^ for intracranial volume, and 1 in Sprooten, et al. ^63^ for global FA (**Supplementary Table 14**). When the significance threshold was relaxed to 5 × 10^−8^, we tagged variants reported in more previous studies, such as 2 in Shen, et al. ^64^ for brain imaging measurements, 2 in Chung, et al. ^65^ for hippocampal volume in dementia, 1 in Chen, et al. ^66^ for putamen volume, and 1 in Christopher, et al. ^67^ for posterior cingulate cortex (**Supplementary Table 15**). For example, we observed colocalizations in region 5q14.3 with previously reported variants for WMH volume and white matter integrity^26^, in 10q26.13 with hippocampal volumes^55^, in 17q21.31 with subcortical^59^ and intracranial^62^ volumes, and in 17q25.1 with WMH volume^26^/burden^28,56^ (**Supplementary Fig. 7**).

Moreover, we found lots of previous associations with other complex traits in different domains (**Supplementary Table 16**). We highlighted 190 variants with psychological traits (e.g., neuroticism^68^, well-being spectrum^69^, general risk tolerance^70^), 179 with cognitive/educational traits (e.g., cognitive ability^71^, educational attainment^72^), 99 with psychiatric disorders (e.g., schizophrenia^73^, MDD^74^, bipolar disorder^75^, ADHD^76^, autism spectrum disorder^77^), 95 with anthropometric traits (e.g., height^78^, body mass index (BMI)^53^), 68 with bone mineral density^79,80^, 54 with smoking/drinking (e.g., smoking^81^, alcohol use disorder^82^), 20 with neurological disorders (e.g., corticobasal degeneration^83^, Parkinson’s disease^84^, Alzheimer’s disease^85^, multiple sclerosis^86^), 18 with sleep (e.g., sleep duration^87^, chronotype^88^), 11 with glioma (glioblastoma or non-glioblastoma) tumors^89,90^, and 6 with stroke^91-93^. For example, white matter associated variants colocalized with many risk variants of cognitive/educational traits as well as brain-related disorders in regions 17q21.31, 6p22.1, and 6p22.2 (**Supplementary Fig. 8**). Strong colocalizations were also found in 7p22.3 with anthropometric traits and bone mineral density, in 10p12.31 with smoking/drinking and anthropometric traits, in 9p21.3 with glioma and stroke, and in 8q24.12 with bone mineral density (**Supplementary Fig. 9**).

To further explore these overlaps, we summarized the number of previously reported variants of other traits that can be tagged by any DTI parameters in each white matter tract (**Supplementary Table 17**). We found that variants associated with psychological, cognitive/educational, smoking/drinking traits and neurological and psychiatric disorders were globally linked to many white matter tracts (**Supplementary Fig. 10**). For traits in other domains, the overlaps may have some tract-specific patterns. For example, 3 of the 6 variants associated with stroke were linked to both SFO and ALIC, and the other 3 were found in superior corona radiata (SCR), anterior corona radiata (ACR), genu of corpus callosum (GCC), body of corpus callosum (BCC), EC, posterior limb of internal capsule (PLIC), and posterior limb of internal capsule (RLIC). In addition, 7 of the 11 risk variants of glioma were associated with splenium of corpus callosum (SCC), 12 of the 18 variants reported for sleep were related to PLIC or inferior fronto-occipital fasciculus (IFO), and 26 of the 68 variants associated with bone mineral density were linked to CST. In addition, more than half of the variants tagged by uncinate fasciculus (UNC) and fornix (FX) had been implicated with anthropometric traits. We carried out voxel-wise association analysis for four representative pleiotropic variants (Methods). **Figure 3** illustrated their genomic locations and voxel-wise effect size patterns in spatial brain maps. rs593720 and rs13198474 had strong effects in corpus callosum (GCC, BCC, and SCC), corona radiata (ACR and SCR), and EX, and the two variants widely tagged psychiatric^94^ and neurological^95^ disorders, as well as psychological^96^ and cognitive/educational^97^ traits. On the other hand, rs77126132 highlighted in SCC and BCC was particularly linked to glioma^89^, and rs798510 in SCR, FX, and PLIC was associated with several anthropometric traits^98^.

**Figure 3:**
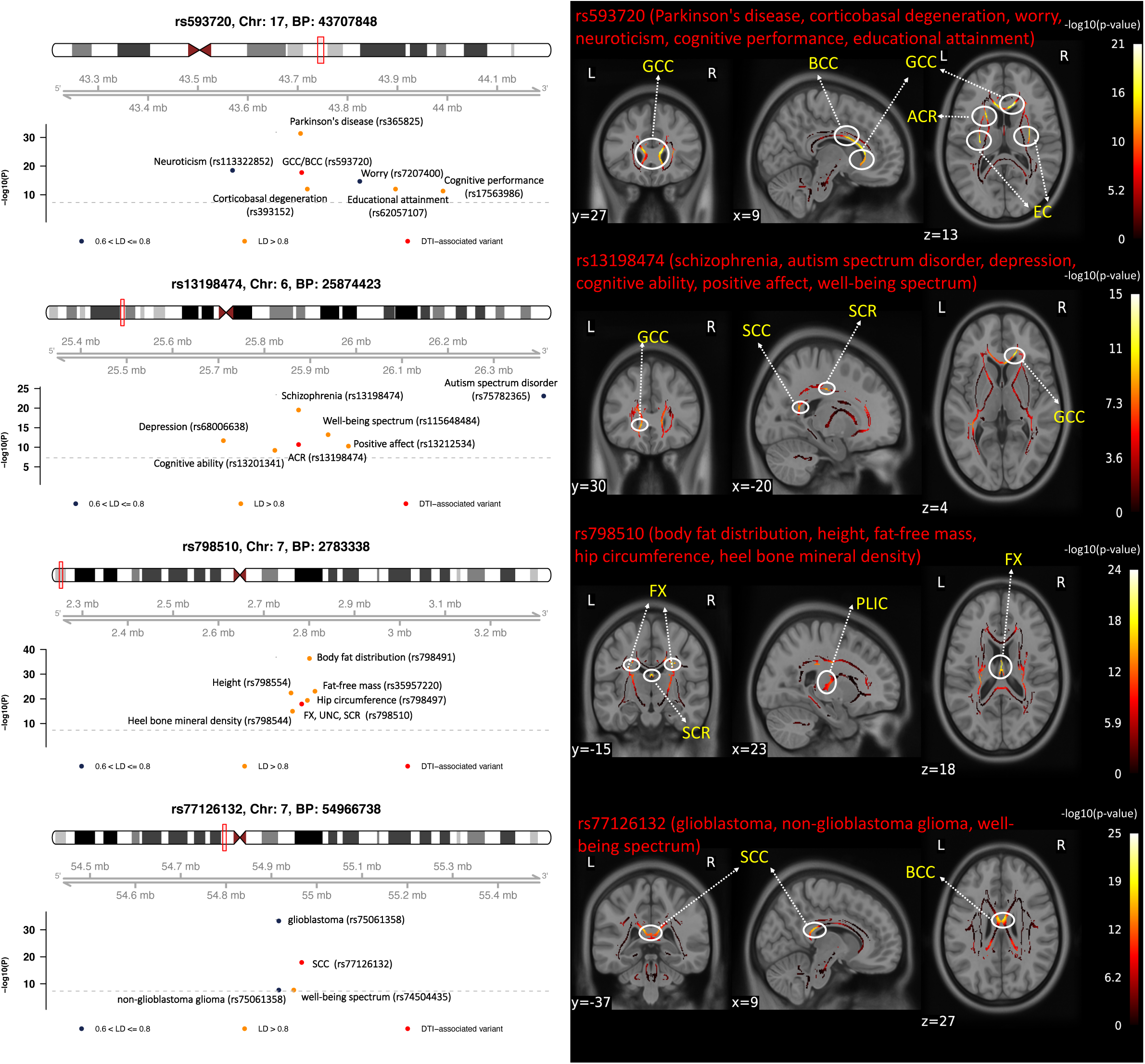
The genomic region and brain spatial map of voxel-wise effect size patterns for four selected pleiotropic variants (n = 33,292 subjects). We labeled previously reported GWAS variants for other complex traits in genomic regions influencing white matter microstructure (left). In spatial maps (right), we illustrate voxel-wise effect sizes of pleiotropic variants in white matter tracts.

### An atlas of genetic correlations with other complex traits

Because of the shared loci associated with both white matter microstructure and other complex traits, we systematically examined their pairwise genetic correlations by using our discovery GWAS summary statistics (*n* = 33,292) and publicly available summary-level data of other 76 complex traits via LDSC (Methods, **Supplementary Table 18**). There were 760 significant pairs between 60 complex traits and 175 DTI parameters after B-H adjustment at 0.05 level (76 × 215 tests, *P* range = (8.6 × 10^−12^, 2.3 × 10^−3^), **Supplementary Table 19**), 38.3% (291/760) of which were detected by PC parameters. We found that DTI parameters were widely correlated with subcortical and WMH volumes (**Supplementary Fig. 11**), brain-related traits (**Supplementary Fig. 12**), and other non-brain traits (**Supplementary Fig. 13**). To validate these results, we performed cross-trait PRS separately on our five European validation GWAS datasets and LDSC on their meta-analyzed summary statistics (*n* = 6,962, Methods). We found that 681 (89.6%) of these 760 significant pairs can be validated in at least one of the six validation analyses after B-H adjustment at 0.05 level (760 tests, *P* range = (1.7 × 10^−10^, 2.9 × 10^−2^), **Supplementary Table 20**), indicating the robustness of our findings. We then reran LDSC after meta-analyzed our UKB British discovery GWAS with these European validation GWAS (*n* = 40,254). The number of significant pairs increased to 855 between 62 complex traits and 178 DTI parameters (**Fig. 4, Supplementary Figs. 14-16** and **Supplementary Table 21**).

**Figure 4:**
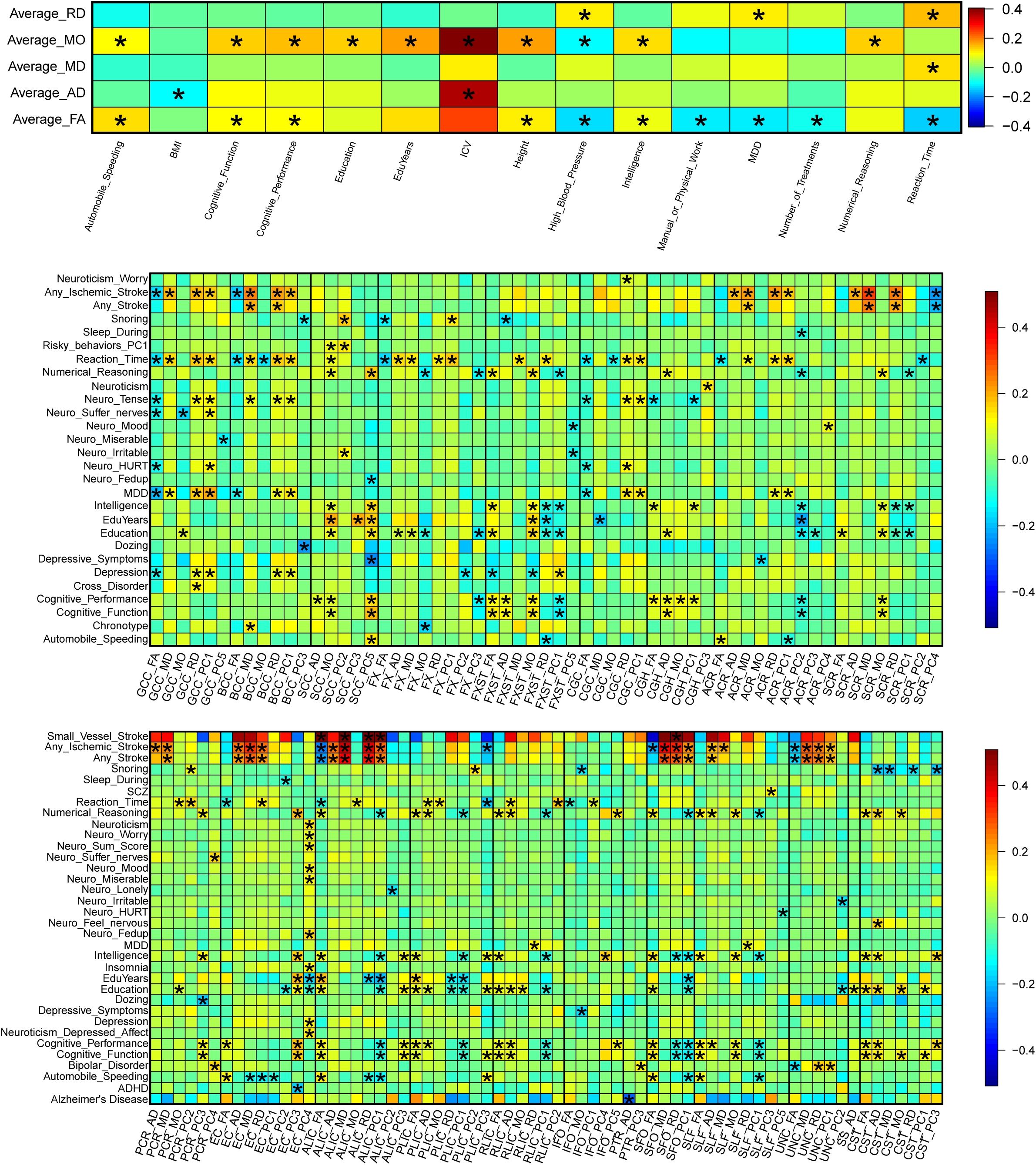
Selected pairwise genetic correlations between white matter microstructure and other complex traits (n = 40,254 subjects). We adjusted for multiple testing by the Benjamini-Hochberg procedure at 0.05 significance level (215 × 76 tests), while significant pairs are labeled with stars. Sample size and detailed information of complex traits can be found in Supplementary Table 18.

We replicated previously reported genetic correlations with cognitive/educational traits^25^, drinking behavior^25^, stroke^23,26^, and MDD^25,26^, and more tract-specific details were revealed. For example, stroke (any subtypes) and ischemic stroke subtypes^92^ (large artery stroke, cardioembolic stroke, and small vessel stroke) showed broad genetic correlations with corpus callosum (GCC and BCC), corona radiata (ACR, SCR, and PCR), limb of internal capsule (PLIC, ALIC), EC, SLF, SFO, and UNC (|gc| range = (0.16, 0.42), *P* < 2.5 × 10^−3^), matching findings in our association lookups. We further observed that small vessel stroke subtype had specific but higher genetic correlations with ALIC and SFO (|gc| range = (0.52, 0.69), *P* < 1.2 × 10^−3^). In contrast, there were no significant genetic correlations detected for large artery and cardioembolic stroke, demonstrating the potentially much stronger genetic links between white matter tracts and small vessel stroke subtype.

More importantly, many new genetic correlations were uncovered for brain-related traits, such as Alzheimer’s disease, ADHD, bipolar disorder, schizophrenia, chronotype, insomnia, neuroticism, and risk tolerance. For example, significant genetic correlation was found between PTR and Alzheimer’s disease (|gc| = 0.30, *P* = 1.7 × 10^−3^), EC and ADHD (|gc| = 0.18, *P* = 4.5 × 10^−5^), UNC and bipolar disorder (|gc| > 0.15, *P* < 4.0 × 10^−4^), and SLF and schizophrenia (|gc| = 0.11, *P* = 2.3 × 10^−3^), matching previously reported case-control differences^12,99-101^ on these tracts. We also found novel significant correlations for non-brain traits, including high blood pressure, height, BMI, bone mineral density, number of non-cancer illnesses and treatments, heavy manual or physical work, smoking, coronary artery disease, lung function, and type 2 diabetes (T2D). For example, high blood pressure was genetically correlated with 19 tracts including SFO, SLF, UNC, EC, and ALIC (|gc| range = (0.09, 0.25), *P* < 2.4 × 10^−3^). Previous research found widespread associations between human brain and these traits, such as bone mineral density^102^, hypertension^103^, T2D^104^, lung function^105^, heart disease^106^, and anthropometric traits^107^. Our findings further illuminate their underlying genetic links. We summarized significant genetic correlations identified in each tract and found that 32.3% (120/372) of these tract-trait genetic correlations can only be detected by PC parameters (**Supplementary Fig. 17** and **Supplementary Table 22**). For example, most of the significant genetic correlations in EC were solely detected by its PC parameters, such as ADHD, BMI, cognitive function, neuroticism, and insomnia.

We explored partial genetic causality among these traits using the latent causal variable^108^ (LCV) model (Methods). As suggested, we conservatively restricted the LCV analysis to pairs with at least nominally significant genetic correlation (*P* < 0.05), significant evidence of genetic causality (B-H adjustment at 0.01 level, 76 × 215 tests), and large genetic causality proportion estimate (|GCP| > 0.6), which were extremely unlikely to be false positives^108^. The LCV model suggested that high blood pressure was partially genetically causal for white matter (|GCP| > 0.67, *P* < 2.2 × 10^−5^, **Supplementary Fig. 18** and **Supplementary Table 23**). On the other hand, white matter may have partially genetically causal effects on insomnia, under sleep, and neuroticism (|GCP| > 0.64, *P* < 7.1 × 10^−8^). These findings may lead to plausible biological hypotheses in future research and suggest the existence of different biological mechanisms underlying the atlas of genetic correlations. More efforts are required to explore causal relationships and the shared biological processes^109^ among these genetically correlated traits.

### Gene-level analysis

We carried out MAGMA^110^ gene-based association analysis for the 215 DTI parameters using our discovery GWAS summary statistics (Methods). There were 3,903 significant gene-level associations (*P* < 1.2 × 10^−8^, adjusted for 215 phenotypes) between 620 genes and 179 DTI parameters (**Supplementary Table 24**), 153 of the associated genes can only be discovered by PC parameters. We replicated 99 of 112 MAGMA genes reported in Zhao, et al. ^25^, 8 white matter-associated genes (*SH3PXD2A, NBEAL1, C1QL1, COL4A2, TRIM47, TRIM65, UNC13D, FBF1*) in Verhaaren, et al. ^56^, 4 (*VCAN, TRIM47, XRCC4, HAPLN1*) in Rutten-Jacobs, et al. ^26^, 3 (*ALDH2, PLEKHG1, TRIM65*) in Traylor, et al. ^27^, 3 (*ALDH2, PLEKHG1, TRIM65*) in Hofer, et al. ^111^, 2 (*TRIM47, TRIM65*) in Fornage, et al. ^28^, and 2 (*GNA12, GNA13*) in Sprooten, et al. ^112^. Most of the other genes had not been implicated with white matter. Many of our MAGMA genes had been linked to other complex traits (**Supplementary Table 25**), such as 70 genes in Anney, et al. ^94^ for autism spectrum disorder or schizophrenia, 50 in Morris, et al. ^79^ for heel bone mineral density, 38 in Hoffmann, et al. ^113^ for blood pressure variation, 51 in Linnér, et al. ^70^ for risk tolerance, 36 in Rask-Andersen, et al. ^98^ for body fat distribution, and 26 in Hill, et al. ^114^ for neuroticism.

Next, we mapped significant variants (*P* < 2.3 × 10^−10^) to genes according to physical position, expression quantitative trait loci (eQTL) association, and 3D chromatin (Hi-C) interaction via FUMA^44^ (Methods). FUMA yielded 1,189 new associated genes (1,630 in total) that were not discovered in MAGMA analysis (**Supplementary Table 26**), replicating 286 of the 292 FUMA genes identified in Zhao, et al. ^25^ and more other genes in previous studies of white matter, such as *PDCD11*^*56*^, *ACOX1*^*56*^, *CLDN23*^*111*^, *EFEMP1*^*26,27,56*^, and *IRS2*^*111*^. More overlapped genes were also observed between white matter and other traits (**Supplementary Table 27)**. Particularly, 876 FUMA genes were solely mapped by significant Hi-C interactions in brain tissues (**Supplementary Table 28**), demonstrating the power of integrating chromatin interaction profiles in GWAS of white matter.

We then explored the gene-level pleiotropy between white matter and 79 complex traits, including nine neurological and psychiatric disorders^115^ studied in Sey, et al. ^115^ and (other) traits studied in our genetic correlation analysis. For brain-related traits, the associated genes were predicted by the recently developed Hi-C-coupled MAGMA^115^ (H-MAGMA) tool (Methods). Traditional MAGMA^110^ was used for non-brain GWAS. H-MAGMA prioritized 737 significant genes for white matter (*P* < 6.3 × 10^−9^, adjusted for 215 phenotypes and two brain tissue types, **Supplementary Table 29**), and we focused on 329 genes that can be replicated in our meta-analyzed European validation GWAS (*n* = 6,962) at nominal significance level (*P* < 0.05, **Supplementary Table 30**). We found that 298 of these 329 genes were associated with at least one of 57 complex traits (**Supplementary Table 31**). **Supplementary Figure 19** and **Supplementary Table 32** display the number of overlapped genes between 57 complex traits and 21 white matter tracts. Most white matter tracts have many pleiotropic genes with other complex traits, aligning with patterns in association lookups and genetic correlation analysis. For example, schizophrenia had 80 overlapped genes with SLF, 71 with CGC, 68 with EC, and 65 with SCR. Global white matter changes in schizophrenia patients had been observed^101,116,117^. Particularly, 230 white matter H-MAGMA genes had been identified in Sey, et al. ^115^ for nine neurological and psychiatric disorders (**Supplementary Table 33**). *NSF*^*118*^, *GFAP*^*119*^, *TRIM27*^*73*^, *HLA-DRA*^*118,120*^, and *KANSL1*^*77,96*^ were associated with five of these disorders, and another 69 genes were linked to at least three different disorders (**Supplementary Fig. 20**). In summary, our analysis largely expands the overview of gene-level pleiotropy, informing the shared genetic influences between white matter and other complex traits.

### Biological annotations

In order to identify tissues and cell types where genetic variation leads to changes in white matter microstructure, we performed partitioned heritability analyses^121^ from the GWAS of global FA and MD within tissue type and cell type specific regulatory elements. First, we utilized regulatory elements across multiple adult and fetal tissues^122^. As expected, both FA and MD had the most significant enrichment of heritability in active gene regulation regions of brain tissues (**Fig. 5a, Supplementary Fig. 21**, and **Supplementary Table 34**). To identify gross cell types, we again performed partitioned heritability using chromatin accessibility data of two brain cell types, neurons (NeuN+) and glia (NeuN-) sampled from 14 brain regions, including both cortical and subcortical^123^. For all regions, we found that significant enrichment of FA and MD heritability existed in glial but not neuronal regulatory elements after B-H adjustment at 0.05 level (**Fig. 5b**). These results are expected as white matter is largely composed of glial cell types. For further resolution on cell types, we tested partitioned heritability enrichment within differentially accessible chromatin of glial cell subtypes, oligodendrocyte (NeuN-/Sox10+), microglia and astrocyte (NeuN-/Sox10-) and two neuronal cell subtypes GABAergic (NeuN+/Sox6+) and glutamatergic neurons (NeuN+/Sox6-) (Methods). Heritability of FA and MD was significantly enriched in oligodendrocyte, microglia, and astrocyte annotations (*P* < 4.8 × 10^−3^). The oligodendrocyte annotation accounted for 10.4% (standard error = 2.6%, *P* = 9.5 × 10^−5^) of the FA heritability while only composed 0.3% of the variants. In contrast, no significant enrichment was observed in neurons (**Fig. 5c**). These analyses imply that common variants associated with white matter microstructure alter the function of regulatory elements in glial cells, particularly oligodendrocytes, the cell type expected to influence white matter microstructure, providing strong support of the biological validity of the genetic associations.

**Figure 5:**
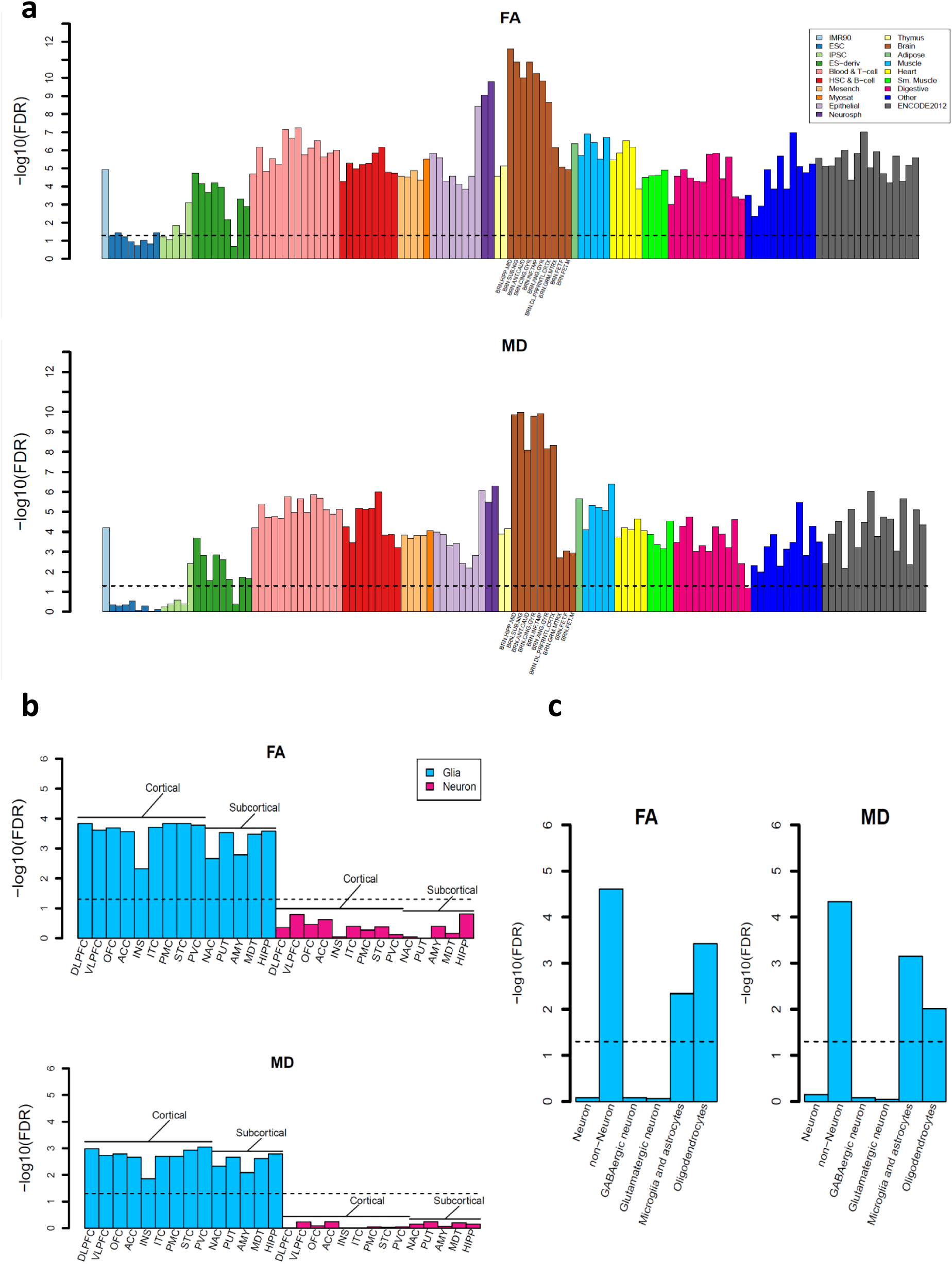
Partitioned heritability enrichment analysis (n = 33,292 subjects). **a)** Heritability enrichment in regulatory elements across tissues and cell types. Brain tissues are labelled in x-axis. **b)** Heritability enrichment in regulatory elements of two brain cell types (neuron and glia) sampled from 14 brain regions. **c)** Heritability enrichment in regulatory elements of glial cell subtypes (non-neuron, including oligodendrocyte and microglia & astrocyte) and neuronal cell subtypes (neuron, including GABAergic and glutamatergic neurons).

To gain more insights into biological mechanisms, we performed several analyses to explore biological interpretations of white matter associated genes. First, MAGMA gene property^110^ analysis was carried out for 13 GTEx^124^ (v8) brain tissues to examine whether the tissue-specific gene expression levels were related to significance between genes and DTI parameters (Methods). After Bonferroni adjustment (13 × 215 tests), we detected 57 significant associations for gene expression in brain cerebellar hemisphere and cerebellum tissues (*P* < 1.8 × 10^−5^, **Supplementary Fig. 22** and **Supplementary Table 35**), suggesting that genes with higher transcription levels on white matter-presented regions also had stronger genetic associations with DTI parameters. In contrast, no signals were observed on regions primarily dominated by grey matter, such as basal ganglia and cortex. Next, we performed drug target lookups in a recently established drug target network^125^, which included 273 nervous system drugs (ATC code starts with “N”) and 241 targeted genes. We found that 19 white matter associated genes were targets for 104 drugs, 43 of which were anti-psychotics (ATC: N05A, target such as *DRD4*) to manage psychosis like schizophrenia and bipolar, 40 were anti-depressants (ATC: N06A, target such as *SLC6A4*) to treat MDD and other conditions, 14 were anti-Parkinson drugs (ATC: N04B, target such as *HTR2B*), and 14 were anti-convulsants (ATC: N03A, target such as *SCN5A*) used in the treatment of epileptic seizures (**Supplementary Table 36**). In addition, we treated white matter associated genes as an annotation and performed partitioned heritability enrichment analysis^121^ for the other 76 complex traits (Methods). After B-H adjustment at 0.05 level, heritability of 54 complex traits was significantly enriched in regions influencing DTI parameters (**Supplementary Fig. 23** and **Supplementary Table 37**). These results suggest the potential clinical values of the genes identified for white matter microstructure.

MAGMA^110^ competitive gene-set analysis was performed for 15,496 gene sets (5,500 curated gene sets and 9,996 GO terms, Methods). We found 180 significant gene sets after Bonferroni adjustment (15,496 × 215 tests, *P* < 1.5 × 10^−8^, **Supplementary Table 38**). The top five frequently prioritized gene sets were “dacosta uv response via ercc3 dn” (M4500), “dacosta uv response via ercc3 common dn” (M13522), “graessmann apoptosis by doxorubicin dn” (M1105), “gobert oligodendrocyte differentiation dn” (M2369), and “blalock alzheimers disease up” (M12921). M4500 and M13522 are *ERCC3*-associated gene sets related to xeroderma pigmentosum (XP) and trichothiodystrophy (TTD) syndromes, which are genetic disorders caused by a defective nucleotide excision repair system^126,127^. In addition to skin symptoms, patients of XP and TTD often reported various neurological deteriorations and white matter abnormalities, such as intellectual impairment^128^, myelin structures degradation^129^, and diffuse dysmyelination^130^. M1105 regulates the apoptosis of breast cancer cells in response to doxorubicin treatment. Clinical research found that breast cancer chemotherapy like doxorubicin was neurotoxic^131^ and can cause therapy-induced brain structural changes and decline in white matter integrity^132^. M2369 plays a critical role in oligodendrocyte differentiation, which mediates the repair of white matter after damaging events^133^, and M12921 is related to the pathogenesis of Alzheimer’s disease^134^.

Several gene sets of rat sarcoma (Ras) proteins, small GTPases, and rho family GTPases were also prioritized by MAGMA, such as “go regulation of small gtpase mediated signal transduction” (GO: 0051056), “go small gtpase mediated signal transduction” (GO: 0007264), “go re gelation of ras protein signal transduction” (GO: 0046578), “go ras protein signal transduction” (GO: 0007265), and “reactome signaling by rho gtpases” (M501). Ras proteins activity is involved in developmental processes and abnormalities of neural cells in central nervous system^135,136^; small and rho family GTPases play crucial roles in basic cellular processes during the entire neurodevelopment process and are closely connected to several neurological disorders^137-139^. We also observed significant enrichment in pathways related to nervous system, including “go neurogenesis” (GO: 0022008), “go neuron differentiation” (GO: 0030182), “go neuron development” (GO: 0048666), “go regulation of neuron differentiation” (GO: 0045664), and “go regulation of nervous system development” (GO: 0051960). Finally, we applied DEPICT^140^ gene-set enrichment testing for 10,968 pre-constituted gene sets (Methods), 7 of which survived Bonferroni adjustment (10,968 × 215 tests, *P* < 2.1 × 10^−8^), such as two gene sets involved in Ras proteins and small GTPases (GO: 0046578 and GO: 0005083) and another two for vasculature and blood vessel developments (GO: 0001944 and GO: 0001568, **Supplementary Table 39**). More MAGMA enriched gene sets can also be detected by DEPICT when the significance threshold was relaxed to 6.5 × 10^−6^ (i.e., not adjusted for testing 215 phenotypes). In summary, our results provide many insights into the underlying biological processes of white matter, suggesting that DTI measures could be useful in understanding the shared pathophysiological pathways between white matter microstructure and multiple diseases and disorders.

## DISCUSSION

In this study, we analyzed the genetic architecture of brain white matter using dMRI scans of 42,919 subjects collected from five publicly accessible data resources. Through a genome-wide analysis, we identified hundreds of previously unknown variants and genes for white matter microstructural differences. Many previously reported genetic hits were confirmed in our discovery GWAS, and we further validated our discovery GWAS in a few replication cohorts. We evaluated the genetic relationships between white matter and a wide variety of complex traits in association lookups, genetic correlation estimation, and gene-level analysis. A large proportion of our findings were revealed by unconventional tract-specific PC parameters. Bioinformatics analyses found tissue and cell-specific functional enrichments and lots of enriched biological pathways. Together, these results suggest the value of large-scale neuroimaging data integration and the application of tract-specific FPCA in studying the genetics of human brain.

One limitation of the present study is that the majority of publicly available dMRI data are from subjects of European ancestry and our discovery GWAS focused on UKB British individuals. Such GWAS strategy can efficiently avoid false discoveries due to population stratifications and heterogeneities across studies^23,141^, but may raise the question that to what degree the research findings can be generalized and applied to global populations^142,143^. In our analysis, we found that the UKB British-derived PRS were still widely significant in Hispanic, Asian, and Black/African American testing cohorts but had reduced performances, especially in Black/African American cohorts. This may indicate that the genetic architecture of white matter is similar but not the same across different populations. Identifying the cross-population and population-specific components of genetic factors for human brain could be an interesting future topic. As more non-European neuroimaging data become available (e.g., the ongoing CHIMGEN project^144^ in Chinese population), global integration efforts are needed to study the comparative genetic architectures and to explore the multi-ethnic genetics relationships among brain and other human complex traits.

### URLs

Brain Imaging GWAS Summary Statistics, https://github.com/BIG-S2/GWAS;

Brain Imaging Genetics Knowledge Portal, https://bigkp.web.unc.edu/;

UKB Imaging Pipeline, https://git.fmrib.ox.ac.uk/falmagro/UK_biobank_pipeline_v_1;

ENIGMA-DTI Pipeline, http://enigma.ini.usc.edu/protocols/dti-protocols/;

PLINK, https://www.cog-genomics.org/plink2/;

GCTA & fastGWA, http://cnsgenomics.com/software/gcta/;

METAL, https://genome.sph.umich.edu/wiki/METAL;

Michigan Imputation Server, https://imputationserver.sph.umich.edu/;

FUMA, http://fuma.ctglab.nl/;

MGAMA, https://ctg.cncr.nl/software/magma;

H-MAGMA, https://github.com/thewonlab/H-MAGMA;

LDSC, https://github.com/bulik/ldsc/;

LCV, https://github.com/lukejoconnor/LCV/;

DEPICT, https://github.com/perslab/depict;

FINDOR, https://github.com/gkichaev/FINDOR;

SuSiE, https://github.com/stephenslab/susieR;

PolyFun, https://github.com/omerwe/polyfun;

NHGRI-EBI GWAS Catalog, https://www.ebi.ac.uk/gwas/home;

The atlas of GWAS Summary Statistics, http://atlas.ctglab.nl/;

## METHODS

Methods are available in the ***Methods*** section.

*Note: One supplementary information pdf file, one supplementary figure pdf file, and one supplementary table zip file are available*.

## ACKNOWLEDGEMENTS

This research was partially supported by U.S. NIH grants MH086633 (H.Z.), MH116527 (TF.L.), and HD079124 (Y.L.). We thank Sophia Cui, Xiaopeng Zong, and Peter Vandehaar for helpful conversations. We thank the individuals represented in the UK Biobank, ABCD, HCP, PING, and PNC studies for their participation and the research teams for their work in collecting, processing and disseminating these datasets for analysis. We gratefully acknowledge all the studies and databases that made GWAS summary data available. This research has been conducted using the UK Biobank resource (application number 22783), subject to a data transfer agreement. Part of the data collection and sharing for this project was funded by the Pediatric Imaging, Neurocognition and Genetics Study (PING) (U.S. National Institutes of Health Grant RC2DA029475). PING is funded by the National Institute on Drug Abuse and the Eunice Kennedy Shriver National Institute of Child Health & Human Development. PING data are disseminated by the PING Coordinating Center at the Center for Human Development, University of California, San Diego. Support for the collection of the PNC datasets was provided by grant RC2MH089983 awarded to Raquel Gur and RC2MH089924 awarded to Hakon Hakonarson. All PNC subjects were recruited through the Center for Applied Genomics at The Children’s Hospital in Philadelphia. Part of the data used in the preparation of this article were obtained from the Adolescent Brain Cognitive Development (ABCD) Study (https://abcdstudy.org), held in the NIMH Data Archive (NDA). This is a multisite, longitudinal study designed to recruit more than 10,000 children age 9-10 and follow them over 10 years into early adulthood. The ABCD Study is supported by the National Institutes of Health and additional federal partners under award numbers U01DA041022, U01DA041028, U01DA041048, U01DA041089, U01DA041106, U01DA041117, U01DA041120, U01DA041134, U01DA041148, U01DA041156, U01DA041174, U24DA041123, U24DA041147, U01DA041093, and U01DA041025. A full list of supporters is available at https://abcdstudy.org/federal-partners.html. A listing of participating sites and a complete listing of the study investigators can be found at https://abcdstudy.org/scientists/workgroups/. ABCD consortium investigators designed and implemented the study and/or provided data but did not necessarily participate in analysis or writing of this report. This manuscript reflects the views of the authors and may not reflect the opinions or views of the NIH or ABCD consortium investigators. HCP data were provided by the Human Connectome Project, WU-Minn Consortium (Principal Investigators: David Van Essen and Kamil Ugurbil; 1U54MH091657) funded by the 16 NIH Institutes and Centers that support the NIH Blueprint for Neuroscience Research; and by the McDonnell Center for Systems Neuroscience at Washington University.

## AUTHOR CONTRIBUTIONS

B.Z., H.Z., Y.L., and J.L.S. designed the study. B.Z., TF. L, Y.Y., X.W., and TY. L analyzed the data. TF. L, Y.S., Z.Z., Y.Y., X.W., TY. L, and D.X., downloaded the datasets, preprocessed dMRI data, and undertook the quantity controls. P.R., M.E.H., J.B., and J.F.F. analyzed brain cell chromatin accessibility data. B.Z. and H.Z. wrote the manuscript with feedback from all authors.

**CORRESPINDENCE AND REQUESTS FOR MATERIALS** should be addressed to H.Z.

## COMPETETING FINANCIAL INTERESTS

The authors declare no competing financial interests.

## METHODS

### GWAS design and Imaging phenotypes

We analyzed the following GWAS datasets separately: 1) the UKB British discovery GWAS, which used data of individuals of British ancestry^52^ from the UKB study (*n* = 33,292); 2) five validation GWAS performed on individuals of European ancestry: UKB White but Non-British (UKBW, *n* = 1,809), ABCD European (ABCDE, *n* = 3,821), HCP (*n* = 334), PING (*n* = 461), and PNC (*n* = 537); 3) two non-European UKB validation GWAS: UKB Asian (UKBA, *n* = 419) and UKB Black (UKBBL, *n* = 211); 4) two non-European non-UKB validation GWAS, including ABCD Hispanic (ABCDH, *n* = 768) and ABCD African American (ABCDA, *n* = 1,257); and 5) a UKB British GWAS with subjects not present in previous GWAS^25^ (also removed the relatives of previous GWAS subjects, *n* = 15,214). See **Supplementary Table 40** for a summary of these GWAS and demographic information of study cohorts. The raw dMRI, covariates and genetic data were downloaded from each data resource. We processed the dMRI data locally using consistent procedures via ENIGMA-DTI pipeline^38,39^ to generate 215 mean and PC DTI phenotypes for 21 predefined white matter tracts (**Supplementary Table 41**). A full description of image acquisition and preprocessing, quality controls, ENIGMA-DTI pipeline, white matter tracts, principle component extraction, and formulas of DTI parameters are detailed in **Supplementary Note**. An overview of tract annotation and imaging procedures is shown in **Supplementary Figures 24-26** and a few image examples are given in **Supplementary Figures 27-30**. For each continuous phenotype or covariate variable, we removed values greater than five times the median absolute deviation from the median value. The ancestry assignment in UKB was based on self-reported ethnic background (Data-Field 21000), whose accuracy was verified in Bycroft, et al. ^52^ For ABCD, we assigned ancestry by a combination analysis using self-reported ethnicity and ancestry inference results from SNPweights^145^, see **Supplementary Note** for details.

### Association discovery and validation

Genotyping and quality controls are documented in **Supplementary Note**. We estimated the SNP heritability by all autosomal SNPs in UKB British discovery GWAS data using GCTA-GREML analysis^43^. The adjusted covariates included age (at imaging), age-squared, sex, age-sex interaction, age-squared-sex interaction, imaging site, as well as the top 40 genetic principle components (PCs) provided by UKB^52^ (Data-Field 22009). The heritability estimates were tested in one-sided likelihood ratio tests. We performed linear mixed model-based association analysis using fastGWA^146^. The same set of covariates as in GCTA-GREML analysis were adjusted. To replicate previous findings, we also performed another UKB British GWAS with subjects not present in previous GWAS^25^. In addition, GWAS were separately performed on European validation datasets UKBW, ABCDE, HCP, PING, and PNC using Plink^147^. In the five validation GWAS, we adjusted for age, age-squared, sex, age-sex interaction, age-squared-sex interaction, and top ten genetic PCs estimated from genetic variants. We also adjusted for imaging sites in ABCD analysis. The meta-analysis was then performed on these validation datasets using METAL^148^ with the sample-size weighted approach.

We applied a few analyses to support the findings in UKB British discovery GWAS. First, the LDSC^48^ software (version 1.0.0) was used to estimate the pairwise genetic correlation between DTI parameter values in discovery GWAS and the meta-analyzed five European validation GWAS (*n* = 6,962). We used the pre-calculated LD scores provided by LDSC, which were computed using 1000 Genomes European data. We used HapMap3^149^ variants and removed all variants in the major histocompatibility complex (MHC) region. In addition, we performed another meta-analysis for the UKB British discovery GWAS and the five European validation GWAS to check whether the *P*-values became smaller after combining these results. Next, polygenic risk scores (PRS) were created on nine validation datasets using the BLUP effect sizes estimated from GCTA-GREML analysis of UKB British discovery GWAS. We used PLINK to generate risk scores in each testing data by summarizing across genome-wide variants, weighed by their BLUP effect sizes. We tried 17 *P*-value thresholds for variant selection using their marginal *P*-values from fastGWA: 1, 0.8, 0.5, 0.4, 0.3, 0.2, 0.1, 0.08, 0.05, 0.02, 0.01, 1 × 10^−3^, 1 × 10^−4^, 1 × 10^−5^, 1 × 10^−6^, 1× 10^−7^, and 1 × 10^−8^. Then, we generated 17 polygenic profiles for each phenotype and reported the best prediction power that can be achieved by a single profile. The association between polygenic profile and phenotype was estimated and tested in linear models, adjusting for the effects of age, gender, and top ten genetic PCs. The additional phenotypic variation that can be explained by polygenic profile (i.e., the incremental R-squared) was used to measure the prediction accuracy.

### Genomic risk loci characterization and comparison with previous findings

We defined genomic risk loci by using FUMA (version 1.3.5e). We input the UKB British discovery GWAS summary statistics after reweighting the *P*-values using functional information via FINDOR^53^. Specifically, FUMA first clumped partially independent significant variants, which were variants with a *P*-value smaller than the predefined threshold and independent of other significant variants (LD *r*^2^ < 0.6, default value). FUMA constructed LD blocks for these independent significant variants by tagging all variants in LD (*r*^2^ ≥ 0.6) with at least one independent significant variant and had a MAF ≥ 0.0005. These variants included those from the 1000 Genomes reference panel that may not have been included in the GWAS. Based on these significant variants, independent lead variants were identified as those that were independent from each other (LD *r*^2^ < 0.1). If LD blocks of independent significant variants were closed (<250 kb based on the closest boundary variants of LD blocks), they were merged to a single genomic locus. Thus, each genomic risk locus could contain more than one independent significant variants and lead variants. We performed functionally-informed fine-mapping by using SuSiE^45^ method via PolyFun^46^ framework for risk loci. The summary statistics from UKB British discovery GWAS were used as input. As suggested, we estimated the LD matrix using our training GWAS individuals. To validate previous findings reported in Zhao, et al. ^25^, we estimated the pairwise genetic correlation between DTI parameter values in previous GWAS and the UKB British GWAS with subjects not included in previous GWAS. We also estimated the replication slope^53^ between two groups of standardized effect sizes. We focused on previously reported top (*P* < 1 × 10^−6^) independent SNPs after LD-based clumping (window size 250, LD *r*^2^ = 0.01). Independent significant variants and all their tagged variants were searched by FUMA in the NHGRI-EBI GWAS catalog (version 2019-09-24) to look for previously reported associations (*P* < 9 × 10^−6^) with any traits. In our UKB British discovery GWAS data, we performed voxel-wise association analysis to illustrate spatial maps for several selected pleiotropic variants. The same set of covariates used in the above tract-based GWAS analysis were adjusted in this voxel-wise analysis.

### Genetic correlation estimation and validation

We used LDSC to estimate the pairwise genetic correlation between DTI parameters and other complex traits. The summary statistics of DTI parameters were from the UKB British discovery GWAS and the summary statistics of other traits were collected from publicly accessible data resources listed in **Supplementary Table 18**. To replicate the significant associations, we reran LDSC using the meta-analyzed summary statistics from the five European validation GWAS. In addition, we also constructed PRS for other complex traits on each of the five validation datasets and tested whether the PRS had significant association with DTI parameters. We used the LD-based pruning (window size 50, step 5, LD *r*^2^ = 0.2) procedure to account for the LD structure in this cross-trait PRS analysis. We also applied the 17 GWAS *P*-value thresholds for variants selection and reported the smallest *P*-value observed in validation data. We applied the LCV^108^ (version 2019-03-14) to explore the genetical causal relationships between DTI parameters and other complex traits. We used meta-analyzed GWAS summary statistics and the pre-calculated LD scores provided by LDSC.

### Gene-level analysis

We first performed gene-based association analysis in UKB British discovery GWAS for 18,796 protein-coding genes using MAGMA^110^ (version 1.07). Default MAGMA settings were used with zero window size around each gene. We then carried out FUMA functional annotation and mapping analysis, in which variants were annotated with their biological functionality and then were linked to 35,808 candidate genes by a combination of positional, eQTL, and 3D chromatin interaction mappings. We chose brain-related tissues/cells in all options and used default values for all other parameters. For the detected genes in MAGMA and FUMA, we performed lookups in the NHGRI-EBI GWAS catalog (version 2020-02-08) again to explore their previously reported associations. We also applied H-MAGMA^115^ (version 2019-11-29) to perform Hi-C coupled gene-based association analysis by integrating Hi-C profiles from fetal and adult brain tissues^150,151^.

### Biological annotations

We performed heritability enrichment analysis via partitioned LDSC^121^. Baseline models were included when estimating the enrichment scores for our tissue type and cell type specific annotations. Methods to prepare in-house chromatin data of three glial cell subtypes and two neuronal cell subtypes can be found in the **Supplementary Note**. We performed gene property analysis for the 13 GTEx^124^ v8 brain tissues via MAGMA. Specifically, we tested whether the tissue-specific gene expression levels can be linked to the strength of the gene-trait association. In addition, we treated DTI associated genes in MAGMA, H-MAGMA or FUMA analysis as an annotation and tested whether the heritability of other complex traits was enriched in this DTI annotation. MAGMA and DEPICT (version 1 rel194) were separately used to explore the implicated biological pathways. MAGMA gene-set analysis examined 5,500 curated gene sets and 9,996 Gene Ontology (GO) terms from the Molecular Signatures Database^152^ (MSigDB, version 7.0) and DEPICT tested 10,968 pre-constructed gene sets using GWAS summary statistics with *P-*value < 10^−5^ as input. All other parameters were set as default.

## Code availability

We made use of publicly available software and tools listed in URLs. Other codes used in our analyses are available upon reasonable request.

## Reporting summary

Further information on research design is available in the Nature Research Reporting Summary.

## Data availability

Our GWAS summary statistics have been shared at https://github.com/BIG-S2/GWAS. The individual-level raw data used in this study can be obtained from five publicly accessible data resources: UK Biobank (http://www.ukbiobank.ac.uk/resources/), ABCD (https://abcdstudy.org/), PING (https://www.chd.ucsd.edu/research/ping-study.html), PNC (https://www.med.upenn.edu/bbl/philadelphianeurodevelopmentalcohort.html), and HCP (https://www.humanconnectome.org/). Our results can also be easily browsed through our knowledge portal https://bigkp.web.unc.edu/.

