## Supplementary_Note for "Common genetic variation influencing human white matter microstructure"

June 16, 2020

### Supplementary Note

#### Genotyping and quality controls

We downloaded the imputed genetic variants data from UKB and HCP data resources, respectively. Genotype imputation was performed locally on the PNC and PING datasets via MACH-Admix (Liu et al., 2013). A full description of the imputation procedures in PNC and PING datasets was detailed supplementary information of Zhao et al. (2019). For the genotype imputation on the ABCD study, we first carried out the following quality control procedures before imputation: 1) exclude subjects with more than 10% missing genotypes; 2) exclude variants with minor allele frequency less than 0.001; 3) exclude variants with missing genotype rate larger than 5%; 4) exclude variants that failed the Hardy-Weinberg test at $1 \times 10^{-9}$ level using only self-identified non-Hispanic white population. We then carried out genotype imputation using the Michigan Imputation Server (`https://imputationserver.sph.umich.edu/`; Das et al. (2016)) and 1000 Genomes Phase 3 (Version 5) reference panel (1000-Genomes-Project-Consortium et al., 2015). Imputed SNPs with a $r^2$-value smaller than 0.3 were removed from the imputation output.

We further performed the following genetic variants data quality controls on each dataset: 1) exclude subjects with more than 10% missing genotypes; 2) exclude variants with minor allele frequency less than 0.01; 3) exclude variants with missing genotype rate larger than 10%; 4) exclude variants that failed the Hardy-Weinberg test at $1 \times 10^{-7}$ level; and 5) remove variants with imputation INFO score less than 0.8.

#### Ancestry assignment in ABCD study

For the ABCD study, we assigned the ancestry to subjects by a combination of self-reported ethnic background and ancestry inference results from the SNPweights software (Chen et al., 2013) (`https://www.hsph.harvard.edu/alkes-price/software/`). Specifically, a four-group continental ancestral reference panel was used in SNPweights to quantify subjects' proportions of African, European, East Asian, and Native American ancestry. We assigned European ancestry to a subject if her/his self-reported ethnic background was European, and also the European proportion in SNPweights was larger than 85%. African American ancestry was assigned to self-reported African American who had $> 90\%$ the combination of African and European ancestry in SNPweights, and also had $< 5\%$ Asian ancestry and $< 5\%$ Native American ancestry. For Hispanic, we focused on self-reported Hispanic subjects who had $> 85\%$ the combination of Native American and European ancestry in SNPweights, and also had $< 5\%$ Asian ancestry and $< 10\%$ African ancestry (Duncan et al., 2018).

### Brain cell chromatin accessibility data processing

**Fluorescence activated nuclear sorting (FANS)** 50mg of frozen brain tissue was homogenized in cold lysis buffer (0.32M Sucrose, 5 mM $CaCl_2$, 3 mM Magnesium acetate, 0.1 mM, EDTA, 10mM Tris-HCl, pH8, 1 mM DTT, 0.1% Triton X-100) and filtered through a 40$\mu$m cell strainer. Following filtration, samples were underlaid with sucrose solution (1.8 M Sucrose, 3 mM Magnesium acetate, 1 mM DTT, 10 mM Tris-HCl, pH8) and centrifuged at 24,000 rpm for 1 hour at $4^o$C. Pellets were re-suspended in 500$\mu$l DPBS, BSA (0.1%) and incubated in anti-NeuN antibody (1:1000, Alexa488 conjugated, Millipore Cat #MAB377X), anti- SOX6 (Kozlenkov et al., 2016) and anti-SOX10 (Ernst et al., 2014), as previously described in Kozlenkov et al. (2016). Prior to fluorescence activated nuclear sorting (FANS), DAPI (Thermoscientific) was added to a final concentration of 1$\mu$g/ml. Nuclei from GABAergic neurons (DAPI+ NeuN+ SOX6+), Glutamatergic neurons (DAPI+ NeuN+ SOX6-), oligodendrocytes (DAPI+ NeuN- SOX10+) and microglia&astrocytes (DAPI+ NeuN- SOX10-) were isolated using a FACSAria flow cytometer (BD Biosciences). This was done in four individuals for three different brain regions (anterior cingulate cortex, dorsolateral prefrontal cortex, and primary visual cortex) resulting in a total of 48 samples.

**Generation of ATAC-seq libraries and sequencing** ATAC-seq reactions were performed using an established protocol (Buenrostro et al., 2015) with minor modifications. In brief, 75,000 sorted nuclei were centrifuged at 500 g for 10 min at $4^o$C. Pellets were re-suspended in transposase reaction mix (25 $\mu$L 2x TD Buffer (Illumina Cat #FC-121-1030) 2.5 $\mu$L Tn5 Transposase (Illumina Cat #FC-121-1030) and 22.5 $\mu$L Nuclease Free H2O) on ice. Reactions were incubated at $37^o$C for 30 min and then purified using the MinElute Reaction Cleanup kit (Qiagen Cat #28204), eluting in 10 $\mu$L of buffer EB. Following purification, ATACseq libraries were generated as described previously (Fullard et al., 2018). Finally, libraries were resolved on 2% agarose gels and fragments, ranging in size from 100-1000 bp, were excised and purified (Qiagen Minelute Gel Extraction Kit – Qiagen Cat#28604). Prior to sequencing, library fragment sizes were estimated using Tapestation D5000 ScreenTapes (Agilent technologies Cat# 5067-5588) and were quantified with the Qubit dsDNA HS assay kit (Invitrogen Cat#Q32851) and by quantitative PCR (KAPA Biosystems Cat#KK4873). Libraries were sequenced on Hi-Seq2500 (Illumina) obtaining $2\times50$ paired-end reads.

**Bioinformatical processing** Sequenced reads were processed as previously described (Fullard et al., 2018). One sample failed quality control based on a low final read count, low signal to noise ratio as evidenced by a low fraction of reads in peaks (FRiP), and the sample being an outlier in clustering analyses. This sample was excluded and the remaining 47 samples were kept. In the Bayesian information criterion-based covariate exploration step, FRiP was picked up as a technical covariate resulting in a model where cell and brain type jointly explained the majority of variance in chromatin accessibility and residuals explained 34%. In contrast to the previous study (Fullard et al., 2018), the four cell type design employed here demanded a modification to the analysis of cell specific chromatin. Here, chromatin was assigned to be specific to a given cell subtype (GABAergic, Glutamatergic, oligodendrocytes, or microglia&astrocyte) if it was significantly more accessible in all pairwise comparisons against the other three cell subtypes. For chromatin specific to the two overall cell types (Neuron or non-Neuron), it was taken by comparing one of the constituent cell subtypes to the two cell subtypes of the other overall cell type, then taking the remaining constituent cell subtype and comparing against the two cell subtypes of the other overall subtype, and then finally taking the union of these two sets. That is, Neuron specific open chromatin was defined as: $((\text{glutamatergic} > \text{oligodendrocyte}) \cap (\text{glutamatergic} > \text{microglia\&astrocyte})) \cup ((\text{GABAergic} > \text{oligodendrocyte}) \cap (\text{GABAergic} > \text{microglia\&astrocyte}))$, and similarly for non-Neuronal specific open chromatin. Note that Neuron and non-Neuron specific open chromatin thus is not simply the sum of open chromatin of their two constituent cell subtypes.

#### Image acquisition and preprocessing

This work made use of diffusion magnetic resonance imaging (dMRI) data from five different data resources, which in general had different imaging protocols. Specifically, the image acquisition and preprocessing procedures were detailed in UK Biobank Brain Imaging Documentation (`https://biobank.ctsu.ox.ac.uk/crystal/crystal/docs/brain_mri.pdf`) for the UK Biobank (UKB) study, Casey et al. (2018) for ABCD study, Satterthwaite et al. (2014) for PNC study, Jernigan et al. (2016) for PING study, and Sotiropoulos et al. (2013) for HCP study. Since UKB is our main discovery dataset, below we briefly introduce the image acquisition and preprocessing procedures used in this study.

**UKB image acquisition** Diffusion weighted imaging (DWI) data of the UK Biobank were acquired at the 2*2*2 mm spatial resolution with multiband acceleration factor of three (i.e., three slices were acquired simultaneously) and two b-values (b = 1,000 and 2,000 s/mm2), anterior-to-posterior is the acquisition phase-encoding direction. For each of these b-values, 50 diffusion-encoding directions were acquired, and there were 100 distinct directions in total. The echo time (TE) was 92 ms and the repetition time (TR) was 3600 ms. The diffusion preparation was a standard (mono-polar) Stejskal-Tanner pulse sequence. The shorter echo time (TE = 92 ms) enables a higher signal-to-noise ratio (SNR) than a twice-refocused (bipolar) sequence at the expense of stronger eddy current distortions (Miller et al., 2016). More details can be found in Section 2.8 of `https://biobank.ctsu.ox.ac.uk/crystal/crystal/docs/brain_mri.pdf`.

**UKB image preprocessing** The UKB dMRI data of the first $\sim$ 20,000 subjects (released in 2018) were preprocessed by the UK Biobank brain imaging team (Alfaro-Almagro et al., 2018). The full pipeline can be found in Section 3 of `https://biobank.ctsu.ox.ac.uk/crystal/crystal/docs/brain_mri.pdf`, referred to as UKB preprocessing pipeline in this note. The pipeline can be divided into two parts: fieldmap generation and Eddy correction. The source codes have been shared at `https://git.fmrib.ox.ac.uk/falmagro/UK_biobank_pipeline_v_1`.

In general, the fieldmap generation workflow in the UKB preprocessing pipeline includes the following steps: 1) all b=0 dMRI images with opposite phase-encoding direction (anterior-posterior (AP) and posterior-anterior (PA)) were analysed to identify the highest-quality pair of AP and PA images; 2) the optimal AP/PA pair was concatenated and then fed into Topup (Andersson et al., 2003) (http://fsl.fmrib.ox.ac.uk/fsl/fslwiki/TOPUP) to estimate the B0 fieldmap and associated dMRI echo planar imaging (EPI) distortions; 3) gradient distortion correction (GDC) was applied and the GDC corrected fieldmap and warpfield were generated; 4) the GDC-corrected magnitude image was linearly aligned to the T1 structural image, and the transformed fieldmap was then masked by the T1-brain mask, resulting in the T1-space fieldmap, which was then transformed back to the B0 space by inverting the previous B0 to T1 registration; and 5) finally, the masked fieldmap was warped to the gradient distorted space using the inverted GDC warpfield.

The Eddy correction workflow has the following steps: 1) using the fieldmap generated in the last step of fieldmap generation workflow, data was corrected for eddy currents and head motion, and outlier-slices (individual slices in the 4D data) were corrected using the Eddy tool (Andersson and Sotiropoulos, 2015, 2016) (http://fsl.fmrib.ox.ac.uk/fsl/fslwiki/EDDY); 2) GDC was then applied, resulting in the 4D output GDC-corrected DWI data; 3) by fitting a diffusion tensor imaging (DTI) model (Basser et al., 1994), DTI parameters FA, L1, L2, L3, V1, V2, V3, MD, and MO were generated.

For the other $\sim$ 20,000 UKB dMRI data recently released in 2019, we downloaded the raw DWI dataset in DICOM format and preprocessed the images locally using the codes shared by the UK Biobank brain imaging team. Structural T1 MRI images were not accessible in DICOM format due to recognizable features of human faces, so we modified the fieldmap generation workflow to generate the brain mask without using the structural imaging information (Supplementary Figure 25). The modified workflow has the following steps: 1) all b=0 dMRI images with opposite phase-encoding direction (AP and PA) were analysed to identify the highest-quality pair of AP and PA images; 2) the optimal AP/PA pair was concatenated and then fed into Topup to estimate the B0 fieldmap and associated dMRI EPI distortions; 3) GDC was applied and the GDC-corrected fieldmap and warpfield were generated; 4) we performed skull-striping on the GDC-corrected magnitude image to generate the brain fieldmap, which was then warped to the gradient distorted space using the inverted GDC warpfield. After fieldmaps were generated, the same Eddy correction steps as above were applied (Supplementary Figure 26).

**UKB image preprocessing quality controls** In summary, there were two quality control (QC) steps in UKB image preprocessing procedures. The first quality control (QC) step is documented in Section 3.4 of `https://biobank.ctsu.ox.ac.uk/crystal/crystal/docs/brain_mri.pdf`. Briefly, if the T1 MRI was considered to be unusable, then the corresponding dMRI would be considered unusable. The "unusable" images were believed to have serious issues such as imaging artefacts/problems or very gross pathologies, so that the downstream image processing pipeline can not perform well. In addition, the raw dMRI data that were incompatible would also be removed and were not processed any further.

Second, as mentioned in the above section, the dMRI data were corrected for eddy currents and head motion, and had outlier-slices corrected using the Eddy tool. GDC was also applied with tools developed by the FreeSurfer and HCP teams (`https://github.com/Washington-University/Pipelines`). Details can be found in Section 3.10 of `https://biobank.ctsu.ox.ac.uk/crystal/crystal/docs/brain_mri.pdf`.

### ENIGMA-DTI pipeline

**More quality control steps** After the above image preprocessing procedures, we obtained the FA, L1, L2, L3, V1, V2, V3, MD, and MO image maps from different datasets. Following `http://enigma.ini.usc.edu/protocols/dti-protocols/`, we conducted two more QC steps. The first was based on directional information of the primary eigenvector. We generated such directional information and manually checked whether the gradients in V1 align appropriately with FA. The Supplementary Figure 27 is an example of what should be expected, where the eigenvector directional information fits the white matter tracts well. In contrast, the example shown in Supplementary Figure 28 is considered to be problematic. In addition, we manually examined the registration performance to make sure all FAs were registered to the template correctly. Specifically, we generated the slices of all registered FA images as in Supplementary Figure 29, and manually detected the abnormal subjects similar to the one in Supplementary Figure 30. The subjects whose FA images did not pass the two QC steps were removed.

**Tract-based analysis** We performed consistent standard registration and quality controls based on the ENIGMA-DTI pipeline (Jahanshad et al., 2013; Kochunov et al., 2014) for different datasets. Specifically, we first used linear registration to register each of the FA images at 1*1*1 mm spatial resolution. We then applied nonlinear registration to align the linearly registered FA images to the ENIGMA FA template and masked the registered FA with a template mask. Next, we projected the ENIGMA skeleton onto the registered FA images to skeletonize them and extracted tract-based FA statistics (tract-averaged mean and functional principal components). Finally, the tract-based statistics (tract-averaged mean) of other four types of DTI parameters (AD, MD, MO, and RD) were obtained by transferring the individual images to the FA template space. The full image data preprocessing and analysis steps are listed below.

1. We carried out image preprocessing steps to process the dMRI data, removed problematic raw images, and conducted corrections for eddy currents, head motions and gradient distortions. The maps of FA, L1, L2, L3, V1, V2, V3, MD, MO parameters at the individual space were generated by fitting a diffusion tensor model using the FSL software (`https://fsl.fmrib.ox.ac.uk/fsl/fslwiki`).

2. We performed additional QCs based on the directional information.

3. We used the FA image to generate the FA mask. This step zeroed the end slices and eroded the image a little bit (`https://github.com/pnlbwh/TBSS/blob/master/TUTORIAL.md`). Abnormal values at the boundary of FA images were excluded from the FA image mask. Then all other diffusion maps were masked by the generated FA mask.

4. We linearly registered the FA image to the JHU ICBM-DTI-81 white-matter labels atlas on the MNI-ICBM-152 space. We used the 9 parameter linear registration and used the correlation ratio as cost function. We performed QC based on linear registration quality. The linear transformation were saved.

5. We nonlinearly registered the linearly aligned FA image to the JHU ICBM-DTI-81 white-matter labels atlas. We performed QC on the nonlinear registration quality. We saved the nonlinear transformation.

6. We warped the L1, L2, L3, MD, MO to the JHU ICBM-DTI-81 white-matter labels atlas according to the linear and nonlinear transformations generated from Step 4 and 5.

7. We skeletonized the registered FA images by projecting the ENIGMA skeleton onto these images. More details can be found in Smith et al. (2006).

8. From the registered FA, L1, L2, L3, MD, RD, MO skeletons, we extracted the regional FA, L1, L2, L3, MD, RD, MO for the predefined white matter tracts according to the JHU ICBM-DTI-81 white-matter labels atlas.

#### White matter tracts

The white matter tracts used in the ENIGMA-DTI protocol were originally from the JHU ICBM-DTI-81 white-matter atlas (Mori et al., 2005; Wakana et al., 2007; Hua et al., 2008). The ENIGMA-DTI protocol linearly registered the FA image to the JHU ICBM-DTI-81 white-matter labels atlas on the MNI-ICBM-152 space, where a total of 48 white matter tract labels were created by hand segmentation of a standard-space average of diffusion MRI tensor maps. A detailed introduction to the tracts included in the JHU ICBM-DTI-81 white-matter atlas can be found in Mori et al. (2008) and Oishi et al. (2008). The middle cerebellar peduncle and pontine crossing tracts were excluded in the ENIGMA-DTI protocol, mainly because the two tracts often fell partially or completely out of the field of view and only mean FA and diffusivity measures for non-zero voxels within the FOV would be calculated (Madden et al., 2012). Thus, there were 46 tracts analyzed in the ENIGMA-DTI protocol and in our analysis, we future took the mean for any left and right tract pairs. The left/right tract pairs were then combined as 25 tracts. We dropped three tracts ('IC', 'CC', and 'CR') because they were the combinations of a few other tracts. That is, 'IC=ALIC+PLIC+RLIC', 'CC'='GCC+SCC+BCC', and 'CR=ACR+SCR+PCR'.

The reproducibility of ENIGMA-DTI protocol pipeline was examined in Jahanshad et al. (2013) and Acheson et al. (2017), and this protocol have widespread applications in recent studies (Kelly et al., 2018; Kim et al., 2019; Dennison et al., 2019). The tracts used in ENIGMA-DTI protocol are clinically and biologically meaningful. For example, the corpus callosum (CC) is the largest fiber tract of the brain, comprising over 190 million axons and connecting the two cerebral hemispheres, linking distant regions of the cerebral cortex (McCarthy-Jones et al., 2018; Edwards et al., 2014). The fornix is a C-shaped bundle of nerve fibers in the brain that acts as the major output tract of the hippocampus, and damage to the fornix can cause difficulty in recalling long-term information such as details of past events.

#### Functional principle component analysis

Functional principal component analysis (FPCA) is a popular method to extract low-dimension structures from voxel-wise neuroimaging data. The principal components (PCs) are the directions that can explain the largest variations of the data. Here we calculated a set of PCs for FA parameter within each tract. The detailed PC extraction steps are listed as follows.

1. For all datasets, we smoothed the skeletonized FA maps using Gaussian kernel with $\sigma = 2mm$ by FSL.

2. For UK Biobank data, we centralized the smoothed DTI parameter skeleton first and then generated the first five PCs for each ROI using the eigenvalue decomposition based on its covariance matrix.

3. For other small datasets (PNC, PING, HCP, and ADNI), we projected the smoothed DTI parameter skeleton of each tract onto the UKB-derived tract-specific eigenvector space, and then extracted the corresponding top five PCs.

#### Formulas of DTI parameters

The DTI parameters were calculated through the FSL software. Specifically, given the processed diffusion weighted images, a diffusion tensor model (Basser et al., 1994) was fitted to characterize the diffusion process, which assumed homogeneity and linearity of the diffusion within each image voxel. For the echo time TE, at a specific voxel $v$, the signal of the dMRI $A(TE)$ follows the following formula:

$$\mathrm{Log}[A(TE)/A(0)] = -\sum_{i,j\leq 3} b_{ij} D_{ij}$$

Since the signal $A(TE)$ can be observed at different $3 \times 3$ matrix $(b_{i,j})$ provided by the UKB team, we can obtain an estimate of the matrix $(D_{ij})$ using linear regression. Thus, the three eigenvalues $L1$, $L2$, $L3$ and eigenvectors V1, V2 and V3 of $(D_{ij})$ can be calculated. The three eigenvalues represent the highest, intermediate, and lowest diffusivity at each voxel. The diffusion measures were explicitly defined as:

$$MD = \frac{L1 + L2 + L3}{3}, \qquad AD = L1, \qquad RD = \frac{L2 + L3}{2},$$

and

$$FA = \sqrt{3/2} \cdot \frac{\sqrt{(L1 - MD)^2 + (L2 - MD)^2 + (L3 - MD)^2}}{\sqrt{L1^2 + L2^2 + L3^2}}.$$

In addition, the mode of anisotropy (MO), i.e., the third moment of the tensor, is defined as

$$MO = \frac{L1L2L3}{||D - \mathrm{trace}(D) \cdot I/3||},$$

where $||x|| = \sqrt{\mathrm{trace}(xx^T)}$.

#### PING methods and authors

Part of the data used in the preparation of this article were obtained from the PING study database (http://ping.chd.ucsd.edu/). PING was launched in 2009 by the National Institute on Drug Abuse and the Eunice Kennedy Shriver National Institute of Child Health & Human Development as a two-year project of the American Recovery and Reinvestment Act. The primary goal of PING has been to create a data resource of highly standardized and carefully curated MRI data, comprehensive genotyping data, and developmental and neuropsychological assessments for a large cohort of developing children aged 3 to 20 years. The scientific aim of the project is, by openly sharing these data, to amplify the power and productivity of investigations of healthy and disordered development in children, and to increase understanding of the origins of variation in neurobehavioral phenotypes. For up-to-date information, see http://ping.chd.ucsd.edu/.

Connor McCabe[1], Linda Chang[2], Natacha Akshoomoff[3], Erik Newman[1], Thomas Ernst[2], Peter Van Zijl[4], Joshua Kuperman[5], Sarah Murray[6], Cinnamon Bloss[6], Mark Appelbaum[1], Anthony Gamst[1], Wesley Thompson[3], Hauke Bartsch[5].

[1]UC San Diego, La Jolla, CA 92093, USA. [2]U Hawaii, Honolulu, HI 96822, USA. [3]Department of Psychiatry, University of California, San Diego, La Jolla, California 92093, USA. [4]Kennedy Krieger Institute, Baltimore, MD 21205, USA. [5]Multimodal Imaging Laboratory, Department of Radiology, University of California San Diego, La Jolla, California 92037, USA. [6]Scripps Translational Science Institute, La Jolla, CA 92037, USA.
