## Supplementary_Figures for "Common genetic variation influencing human white matter microstructure"

June 13, 2020

### Supplementary Figures

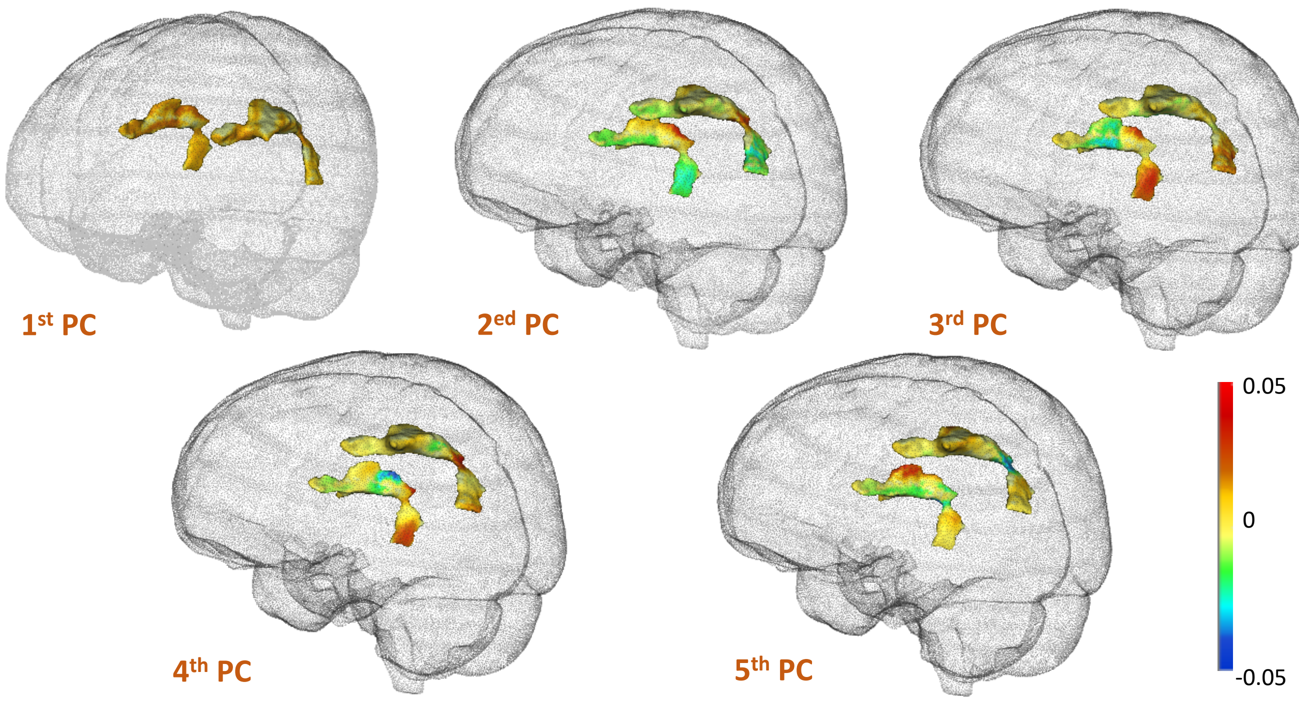


**Supplementary Figure 1:** Illustration of the functional principal component (PC) coefficients for the top five fractional anisotropy (FA) PCs of superior longitudinal fasciculus (SLF).

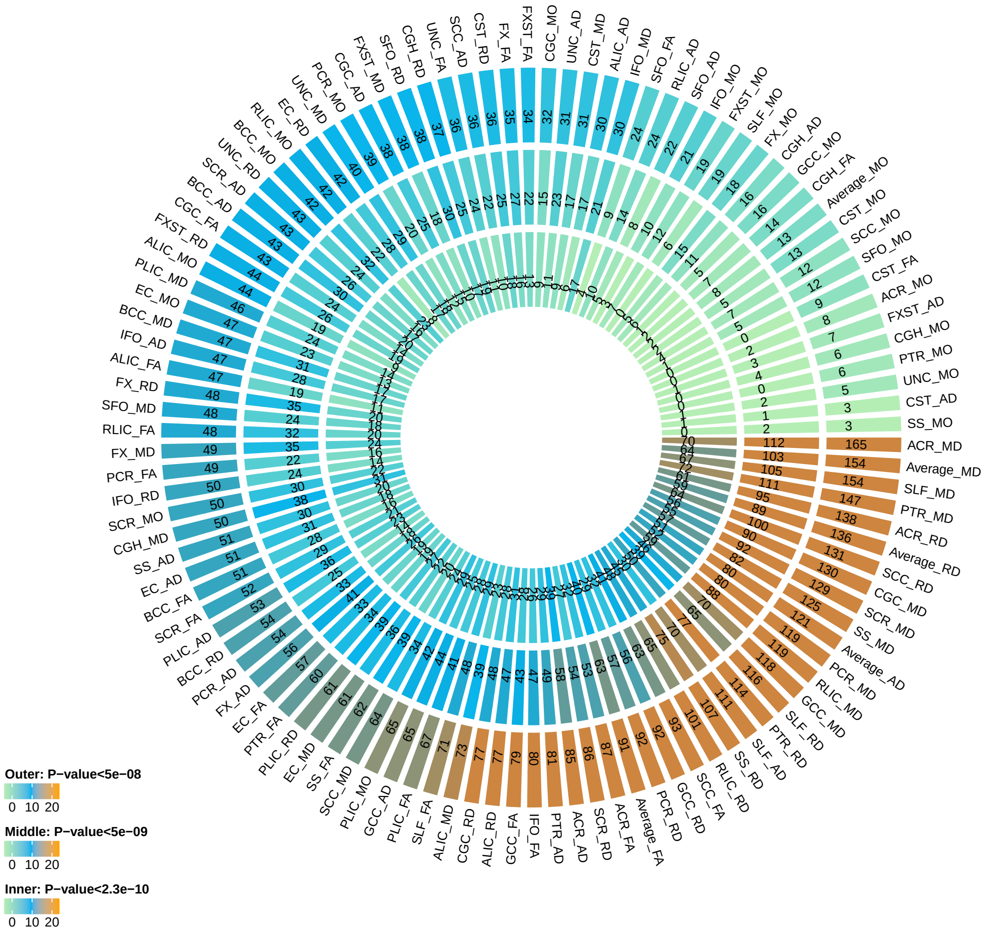


**Supplementary Figure 2:** Number of independent significant variants discovered for 110 mean parameters in UKB British discovery GWAS (n=33,292 subjects) at different significance levels. Outer layer: P-value $< 5 \times 10^{-8}$; middle layer: P-value $< 5 \times 10^{-9}$; and inner layer: P-value $< 2.3 \times 10^{-10}$. The $2.3 \times 10^{-10}$ threshold corresponds to additionally adjusting for testing 215 imaging phenotypes with the Bonferroni correction. The p-values are raw p-values of two-sided test statistics in mixed linear model-based association analysis generated by fastGWA (`https://cnsgenomics.com/software/gcta/`).

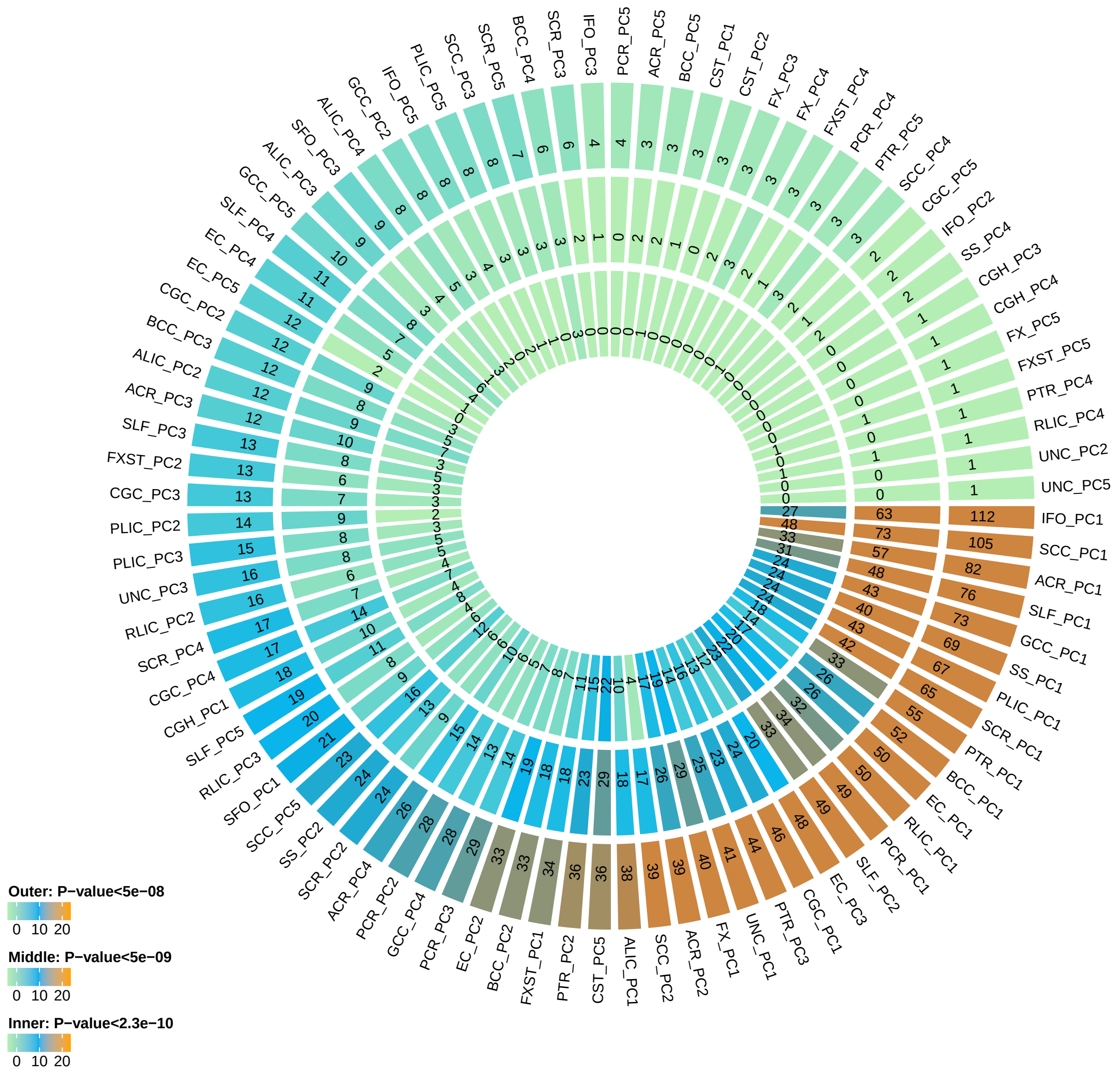


**Supplementary Figure 3:** Number of independent significant variants discovered for 105 PC parameters in UKB British discovery GWAS (n=33,292 subjects) at different significance levels. Outer layer: P-value $< 5 \times 10^{-8}$; middle layer: P-value $< 5 \times 10^{-9}$; and inner layer: P-value $< 2.3 \times 10^{-10}$. The $2.3 \times 10^{-10}$ threshold corresponds to additionally adjusting for testing 215 imaging phenotypes with the Bonferroni correction. The p-values are raw p-values of two-sided test statistics in mixed linear model-based association analysis generated by fastGWA (`https://cnsgenomics.com/software/gcta/`).

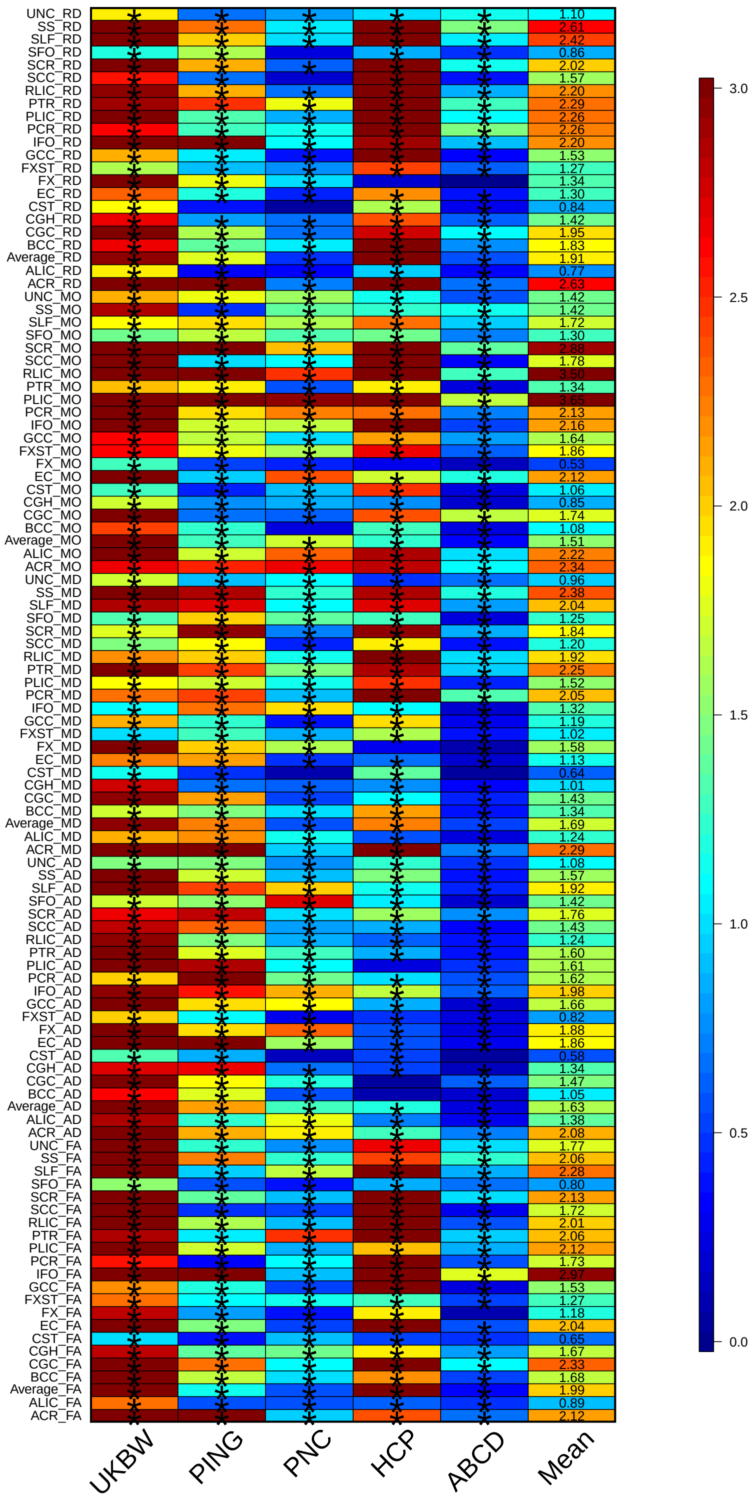

**Supplementary Figure 4:** Prediction accuracy (i.e., incremental R-squared) of polygenic risk scores (PRS) for 110 mean parameters constructed by UKB British discovery GWAS (n=33,292 subjects) summary statistics on the five independent European ancestry datasets and their average across these datasets. Significant PRS after B-H adjustment at 0.05 level were labeled with stars.

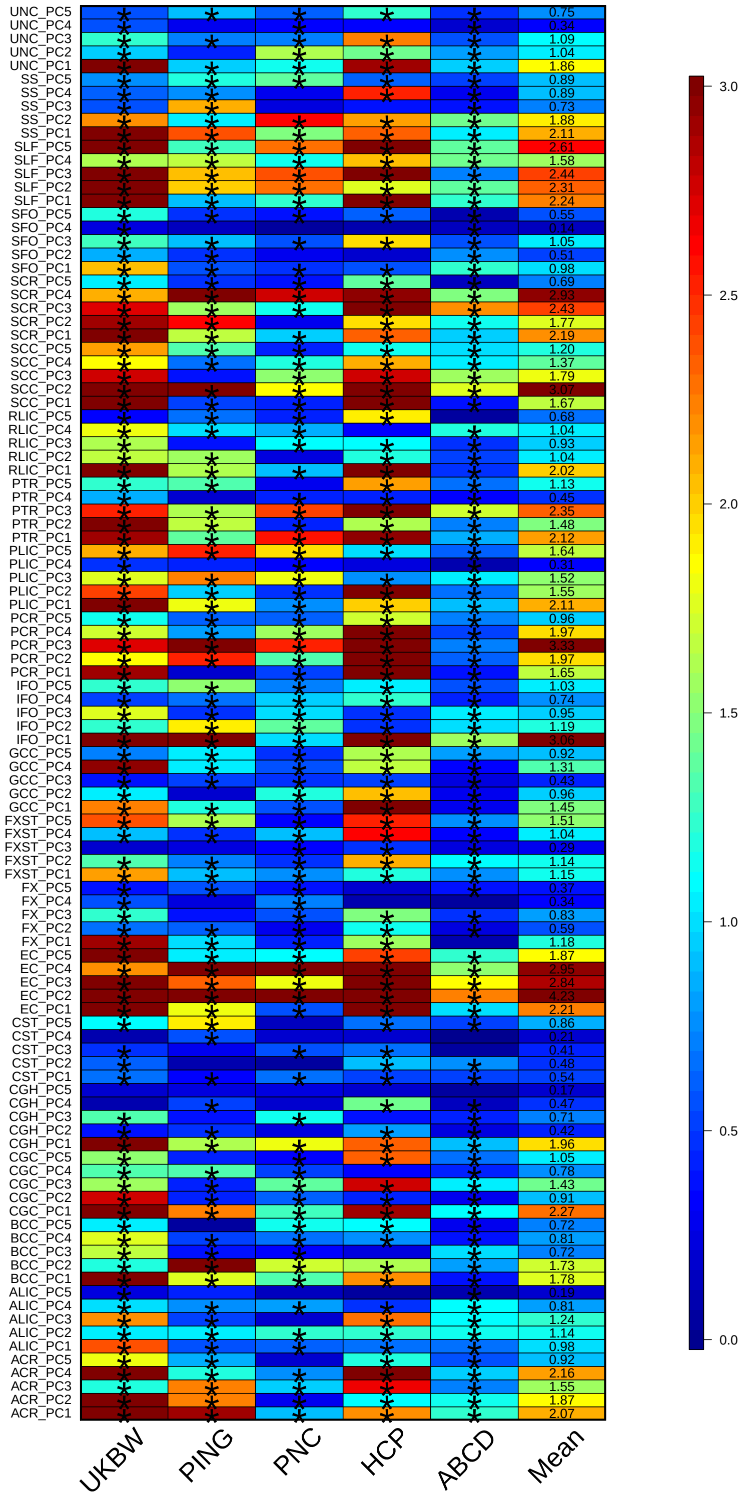


**Supplementary Figure 5:** Prediction accuracy (i.e., incremental R-squared) of polygenic risk scores (PRS) for 105 PC parameters constructed by UKB British discovery GWAS (n=33,292 subjects) summary statistics on the five independent European ancestry datasets and their average across these datasets. Significant PRS after B-H adjustment at 0.05 level were labeled with stars.

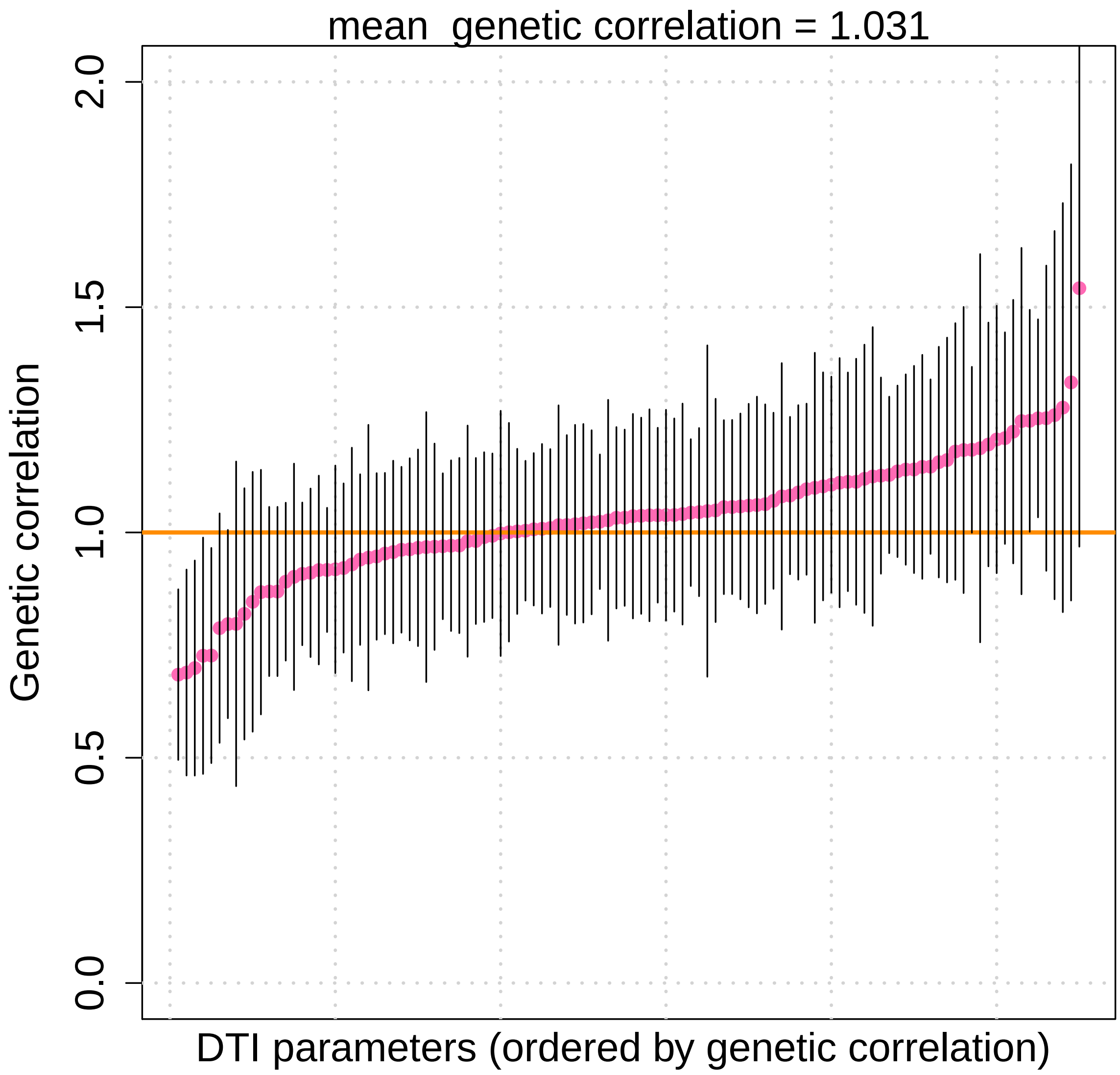


**Supplementary Figure 6:** Genetic correlation estimates and 95% confidence intervals for the 110 mean parameters between previous UKB GWAS (n=17,706 subjects) and the new UKB British validation GWAS (n=15,214 subjects).

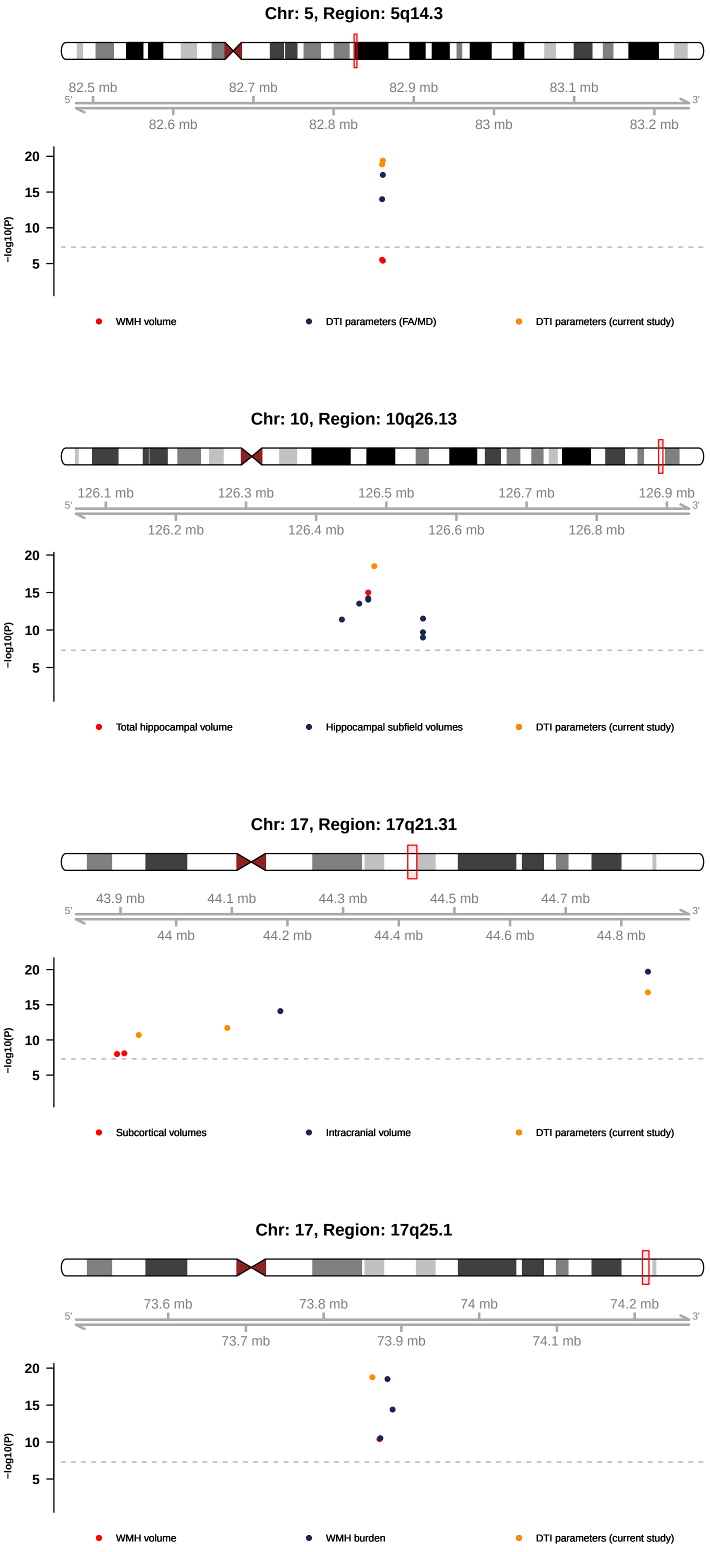


**Supplementary Figure 7:** Colocalizations (LD $r^2 \geq 0.6$) between DTI associated variants in UKB British discovery GWAS (n=33,292 subjects) and previously reported variants for brain imaging traits in selected genome regions.

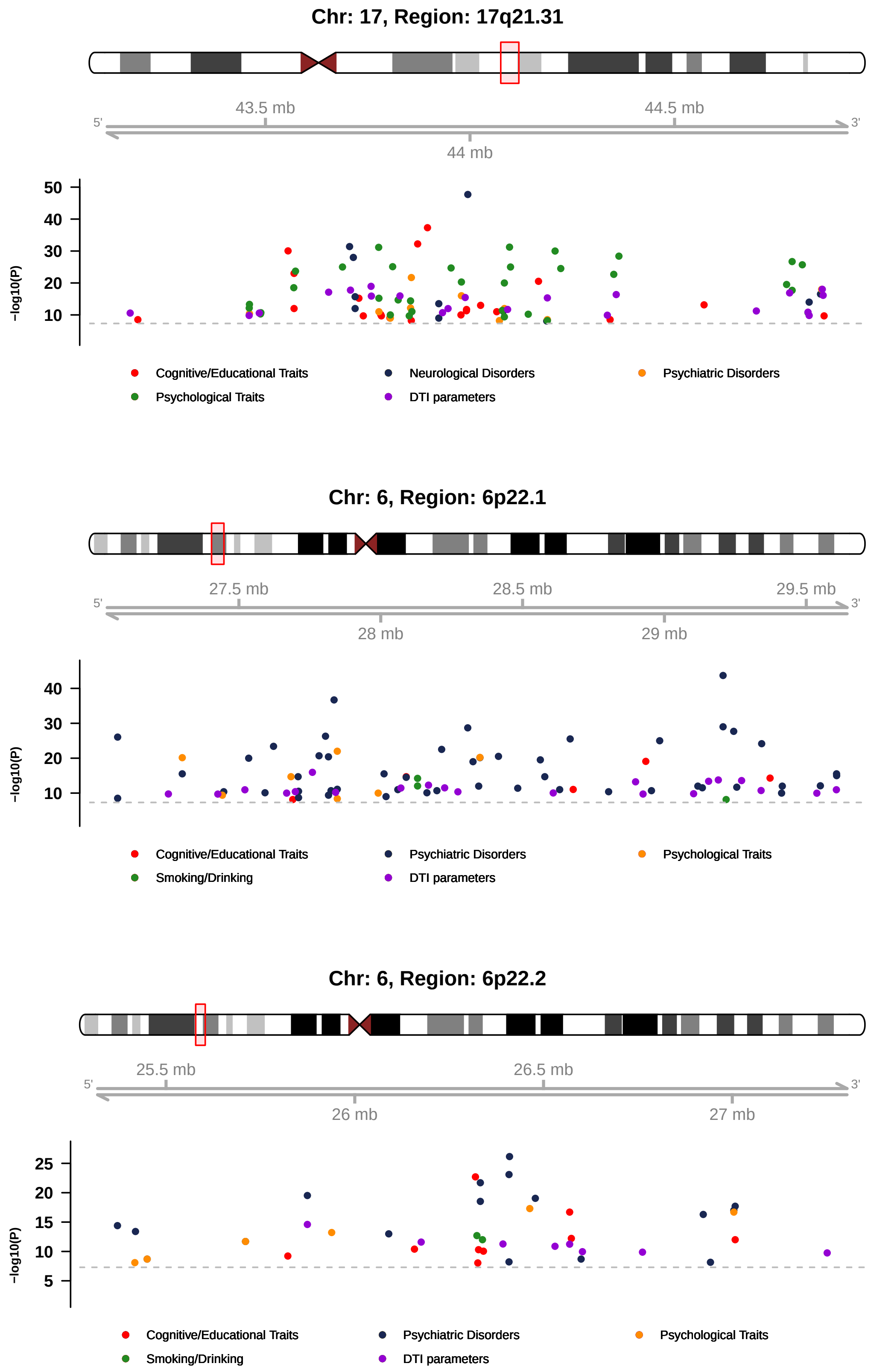


**Supplementary Figure 8:** Colocalizations (LD $r^2 \geq 0.6$) between DTI associated variants in UKB British discovery GWAS (n=33,292 subjects) and previously reported variants for other complex traits in selected genome regions.

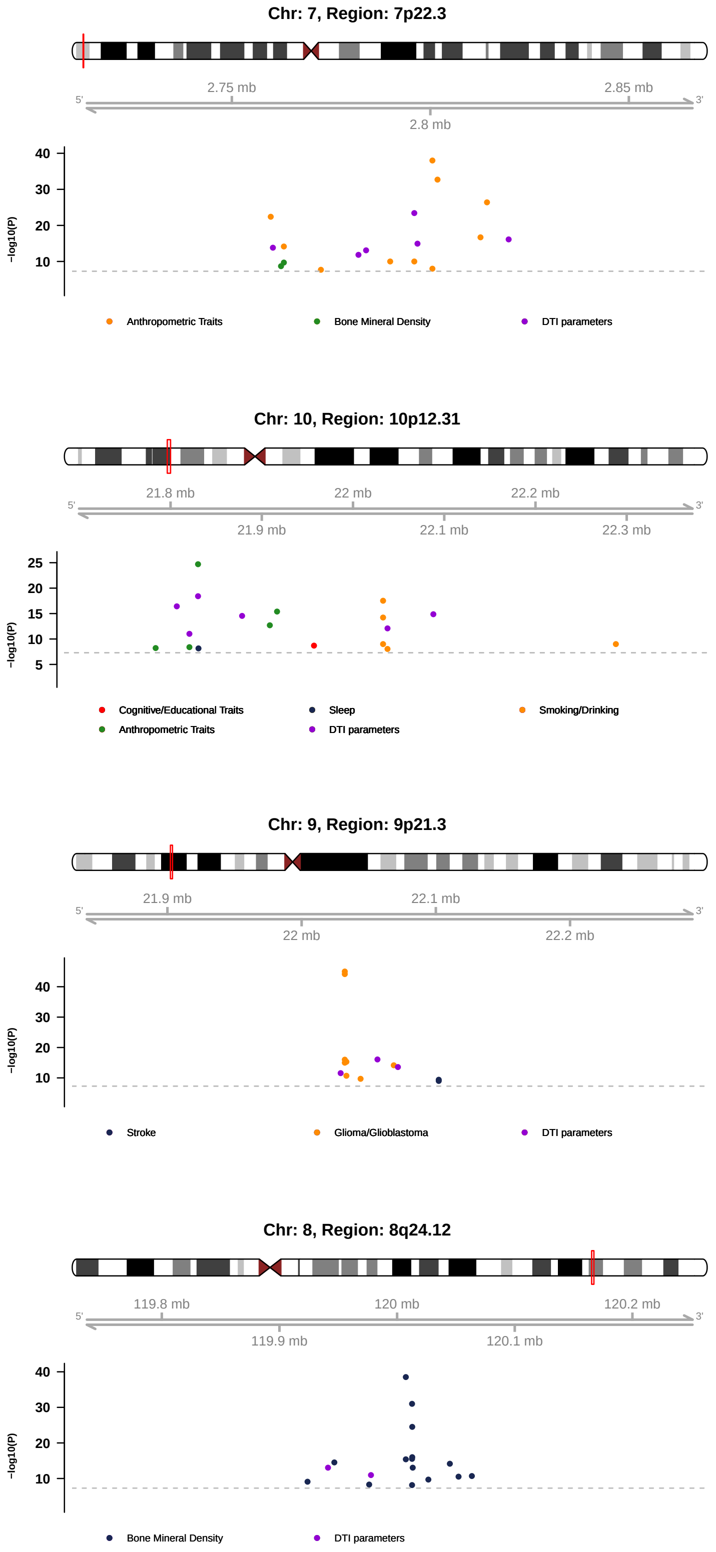


**Supplementary Figure 9:** Colocalizations (LD $r^2 \geq 0.6$) between DTI associated variants in UKB British discovery GWAS (n=33,292 subjects) and previously reported variants for other complex traits in selected genome regions.

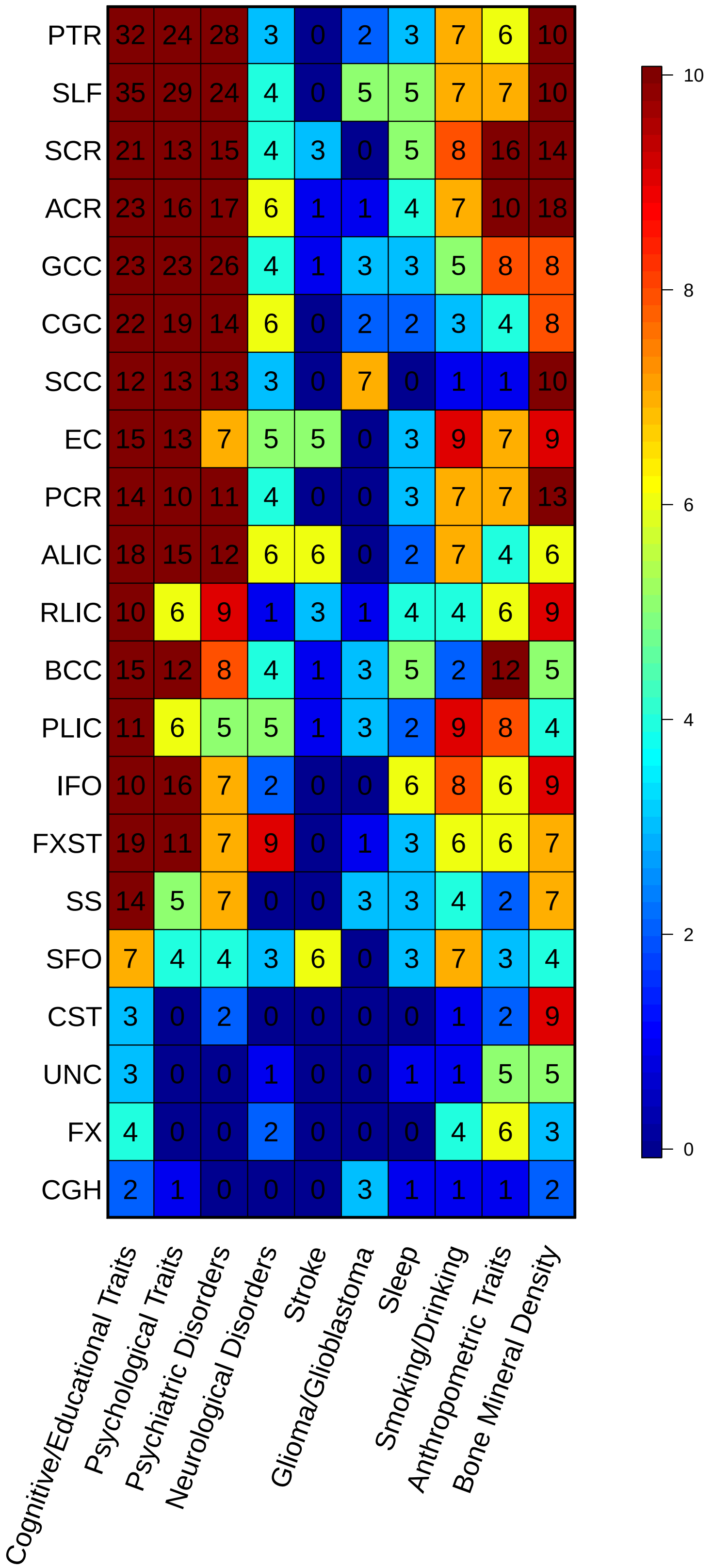


**Supplementary Figure 10:** Number of previously reported GWAS variants for other complex traits that are associated with white matter tracts.

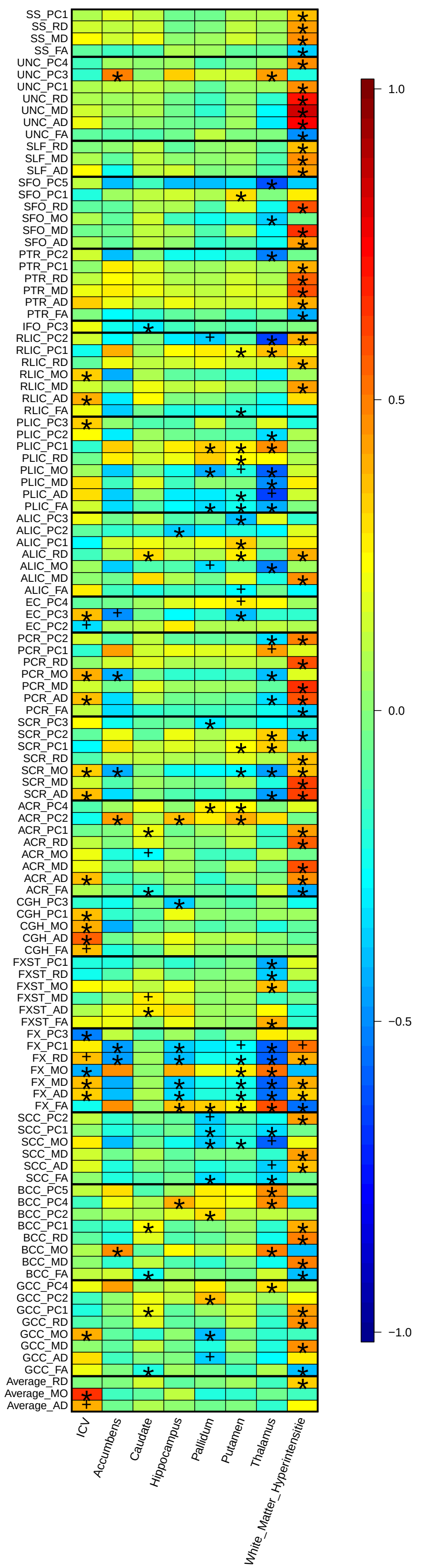

**Supplementary Figure 11:** Genetic correlation estimates between DTI parameters (n=33,292 subjects) and brain volume traits. Stars represent significant associations that can be validated in at least one of the six validation analyses after B-H adjustment at 0.05 level; "+"s represent significant associations that were not validated.

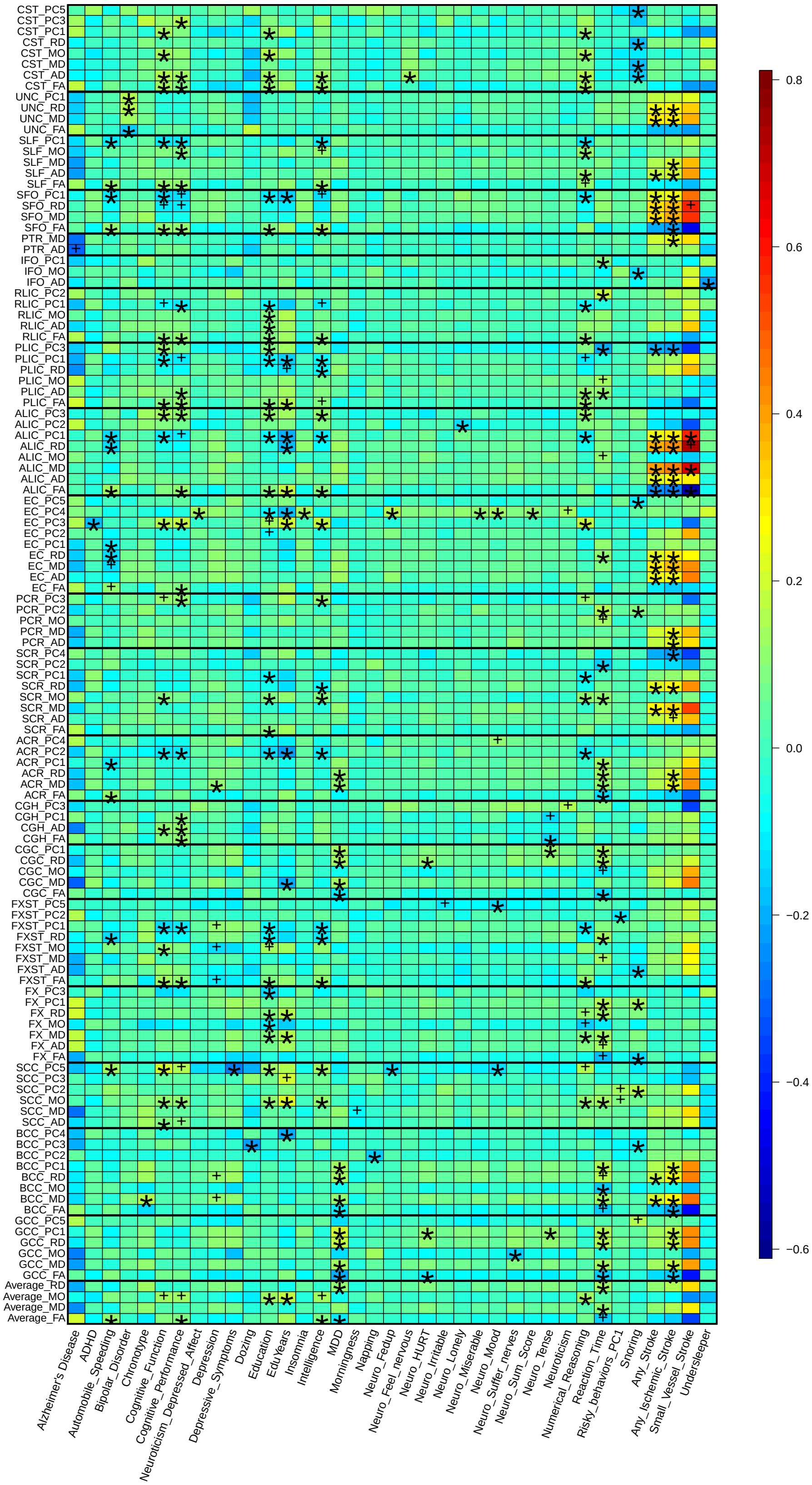


**Supplementary Figure 12:** Genetic correlation estimates between DTI parameters (n=33,292 subjects) and brain-related traits. Stars represent significant associations that can be validated in at least one of the six validation analyses after B-H adjustment at 0.05 level; "+"s represent significant associations that were not validated.

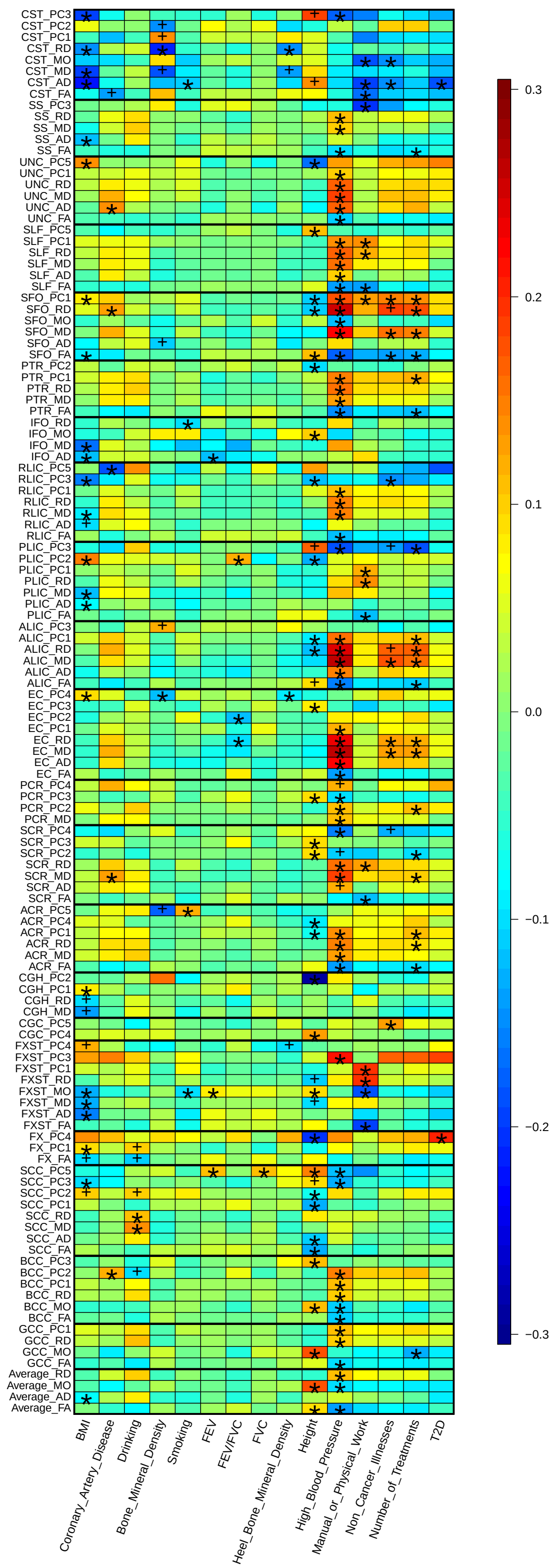


**Supplementary Figure 13:** Genetic correlation estimates between DTI parameters (n=33,292 subjects) and non-brain traits. Stars represent significant associations that can be validated in at least one of the six validation analyses after B-H adjustment at 0.05 level; "+"s represent significant associations that were not validated.

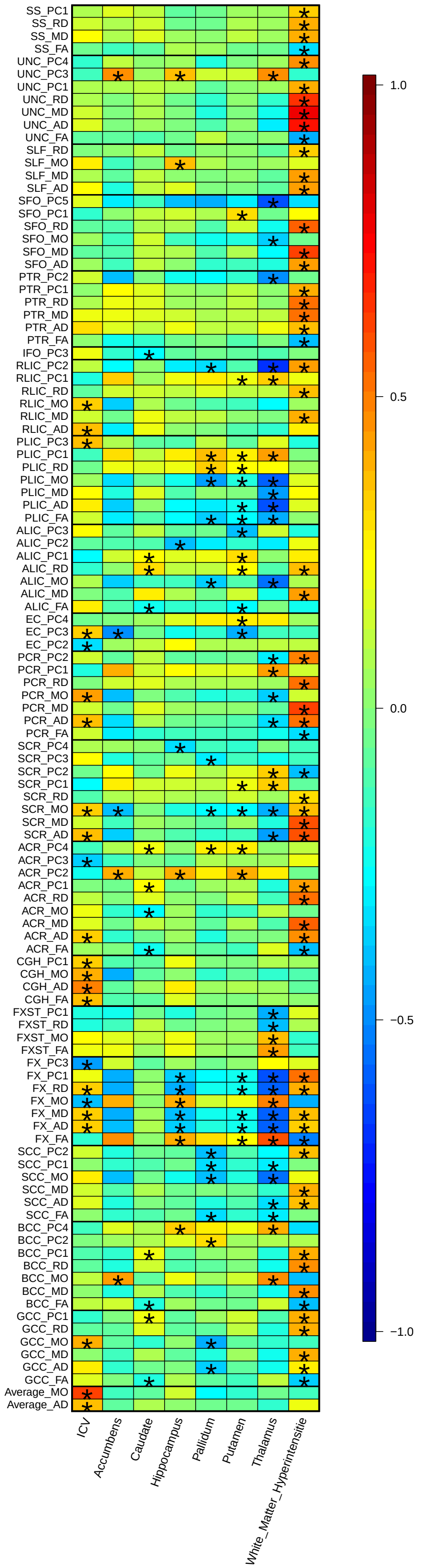


**Supplementary Figure 14:** Genetic correlation estimates between DTI parameters (n=40,254 subjects, meta-analyzed GWAS) and brain volume traits. Stars represent significant associations after B-H adjustment at 0.05 level.

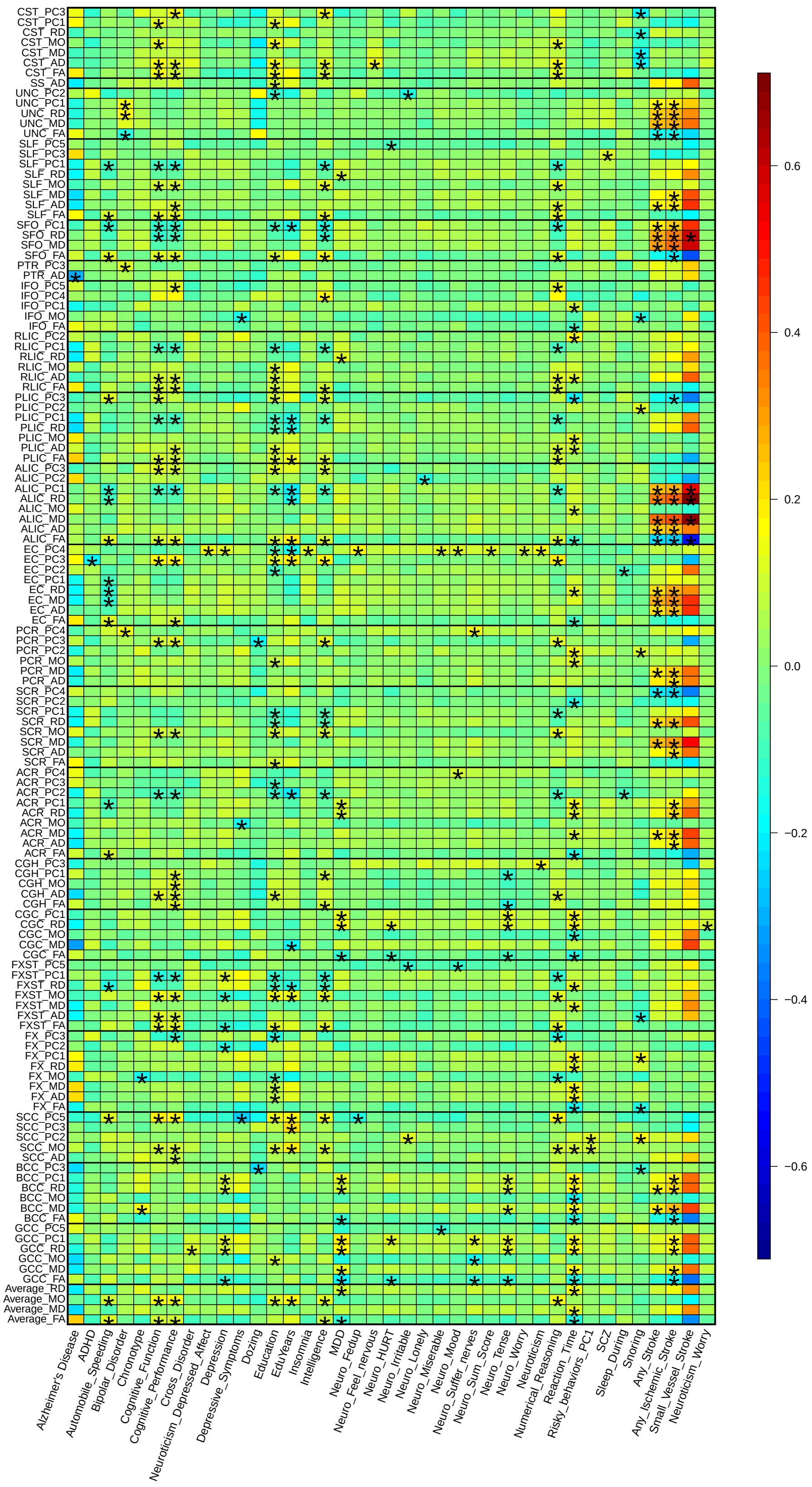

**Supplementary Figure 15:** Genetic correlation estimates between DTI parameters (n=40,254 subjects, meta-analyzed GWAS) and brain-related traits. Stars represent significant associations after B-H adjustment at 0.05 level.

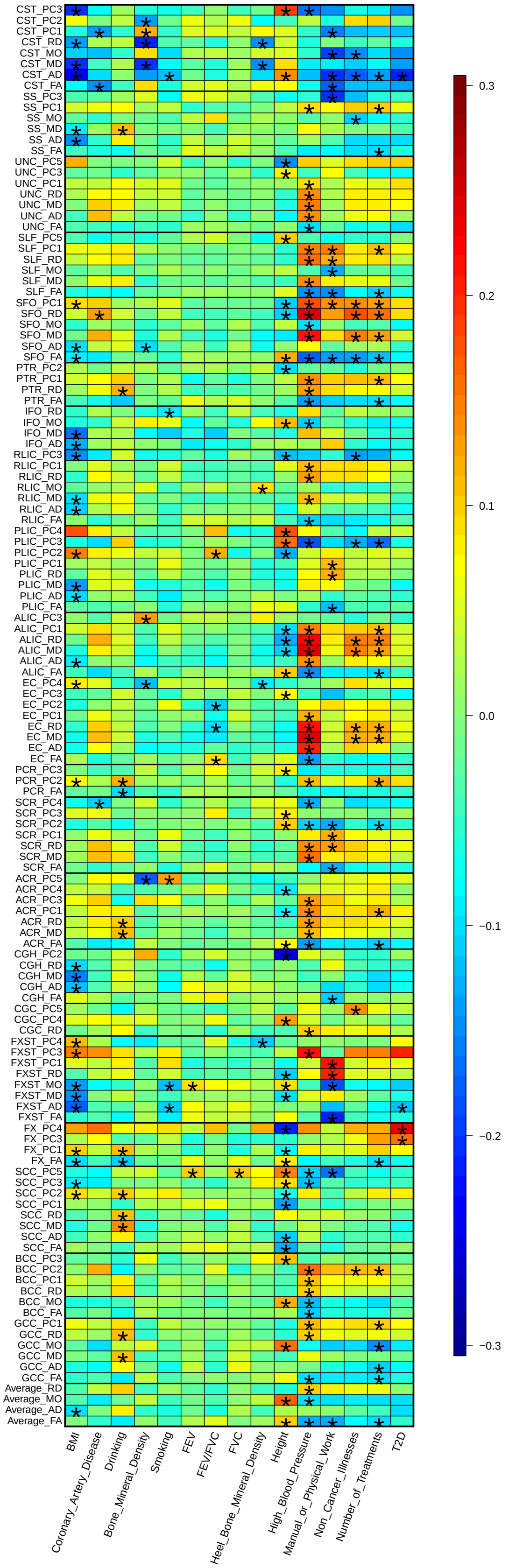


**Supplementary Figure 16:** Genetic correlation estimates between DTI parameters (n=40,254 subjects, meta-analyzed GWAS) and non-brain traits. Stars represent significant associations after B-H adjustment at 0.05 level.

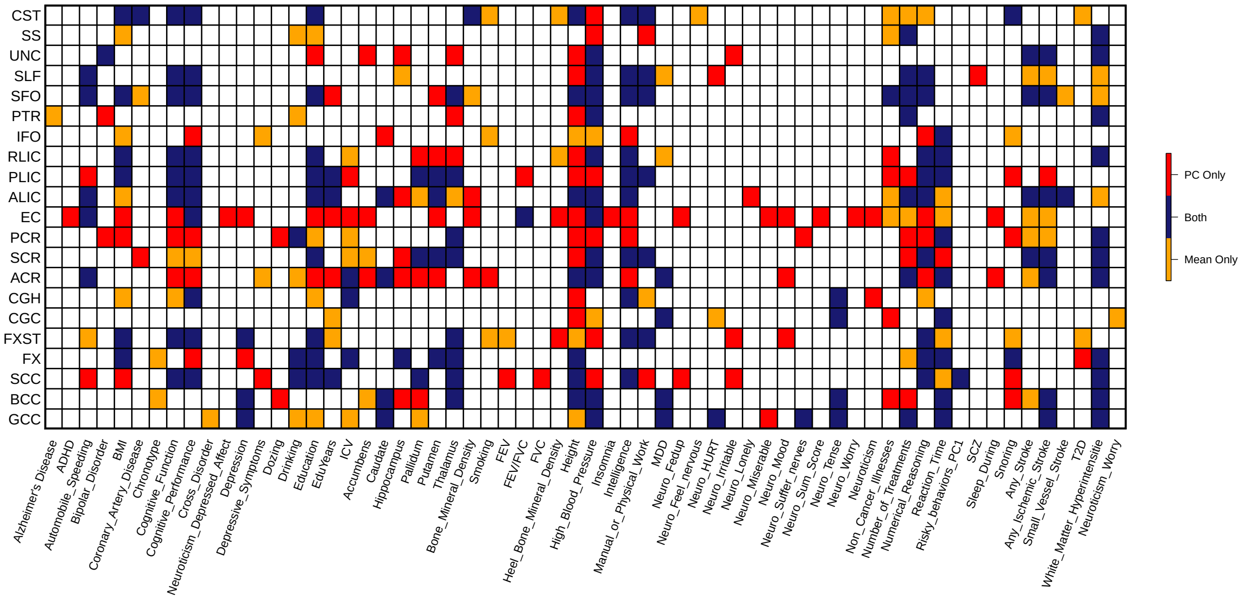


**Supplementary Figure 17:** Significant pairwise genetic correlations between white matter tracts and other complex traits (n=40,254 subjects). The pairs were divided into three groups and labeled with three different colors: "Mean only" (orange) represents pairs solely identified by mean DTI parameters; "PC only" (red) refers to pairs uniquely detected by FA PC DTI parameters; and "Both" (blue) corresponds to pairs that can be found by both mean and PC parameters.

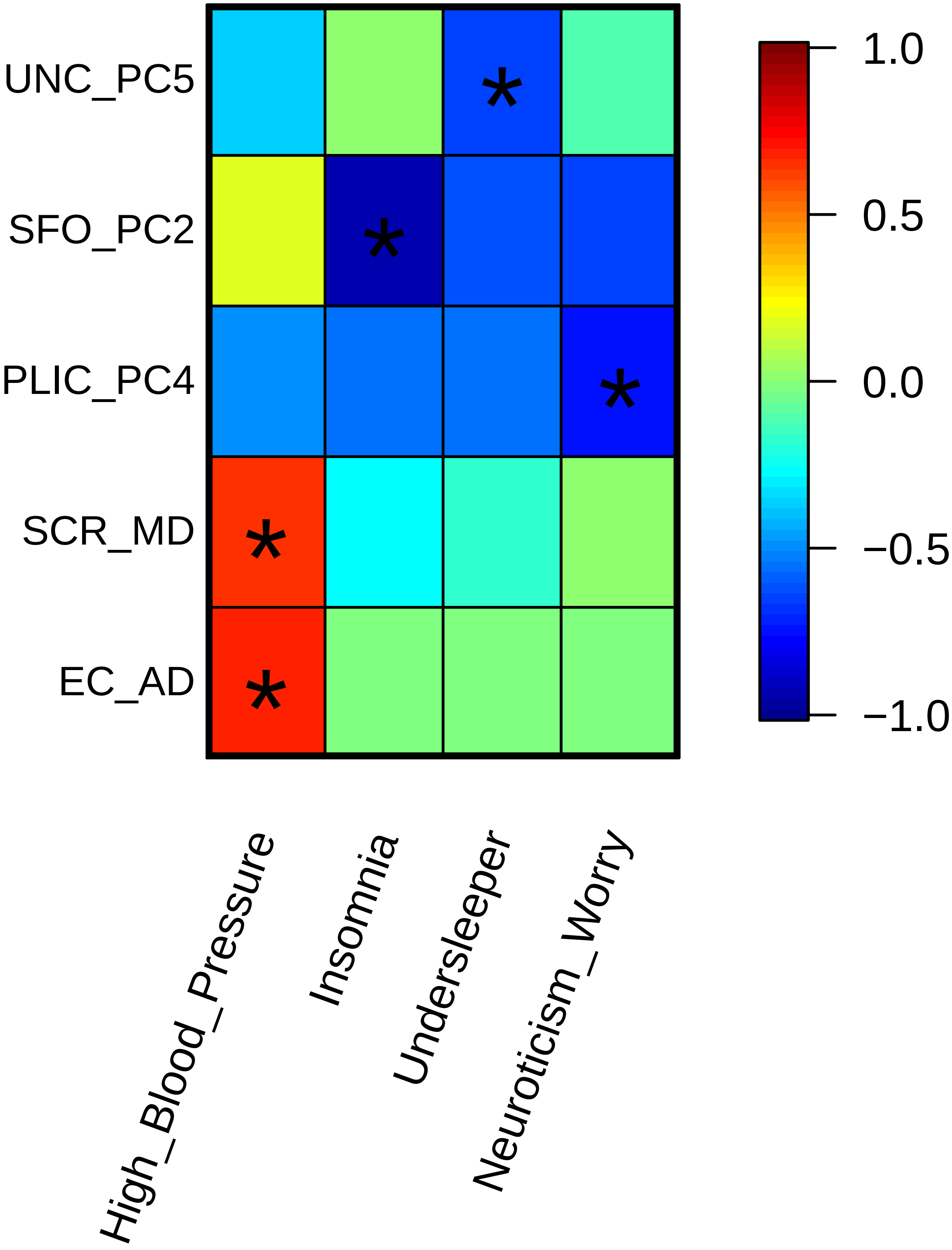


**Supplementary Figure 18:** Genetic causality proportion estimates between DTI parameters (n=40,254 subjects) and other traits by using latent causal variable (LCV) model. Stars represent significant partially genetically causal relationships.

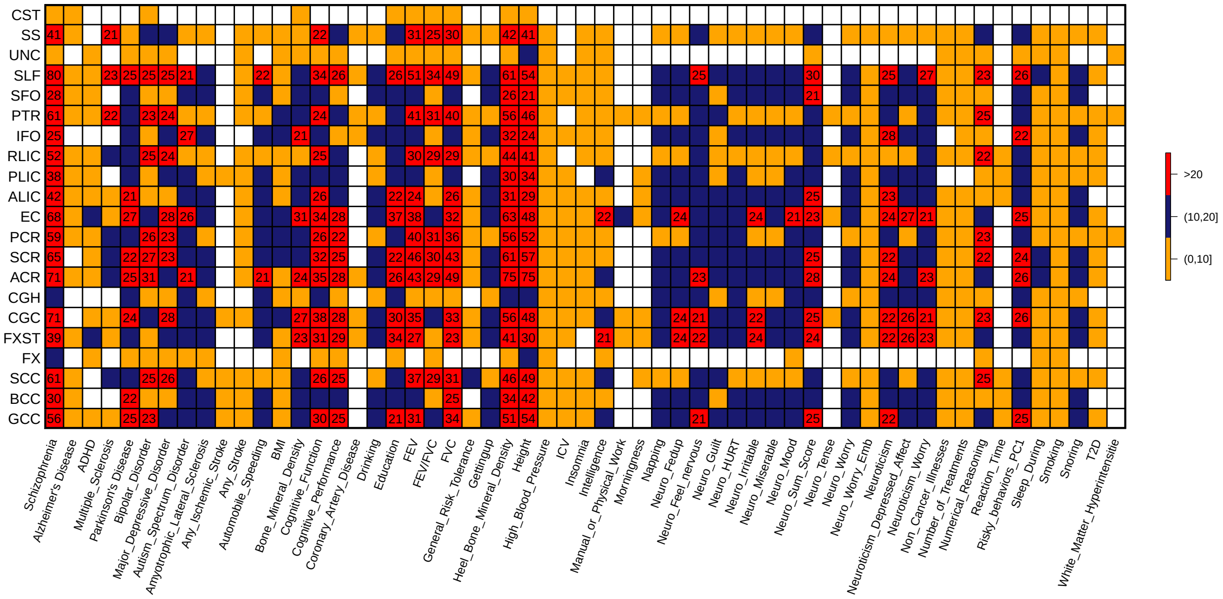


**Supplementary Figure 19:** Number of overlapped genes between 57 complex traits and 21 white matter tracts in gene-based association analysis (n=33,292 subjects). The displayed numbers are the number of overlapped genes between each tract-trait pair in gene-based association analysis.

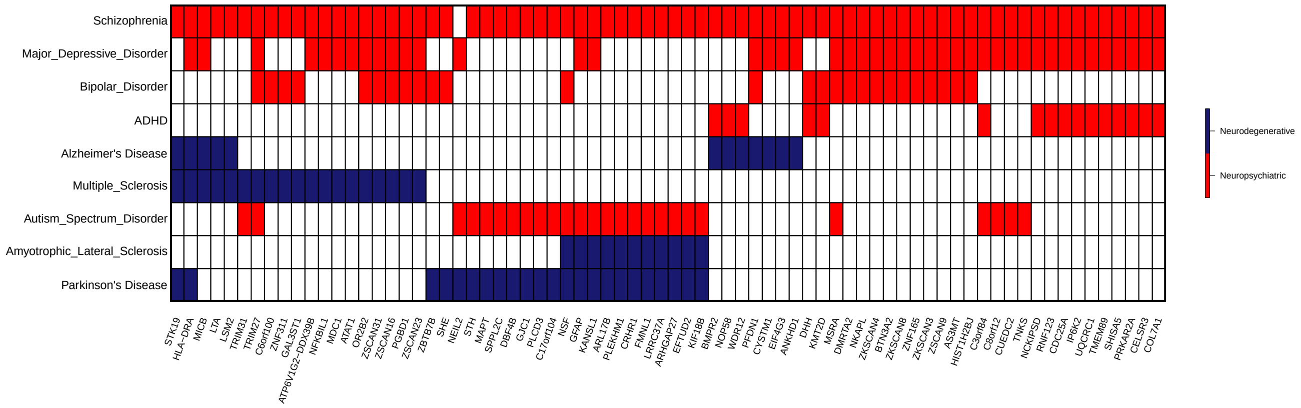


**Supplementary Figure 20:** Pleiotropic genes identified in HMAGMA of DTI parameters (n=33,292 subjects) that have been linked to nine neurological and psychiatric disorders in Sey, et al. (2020).

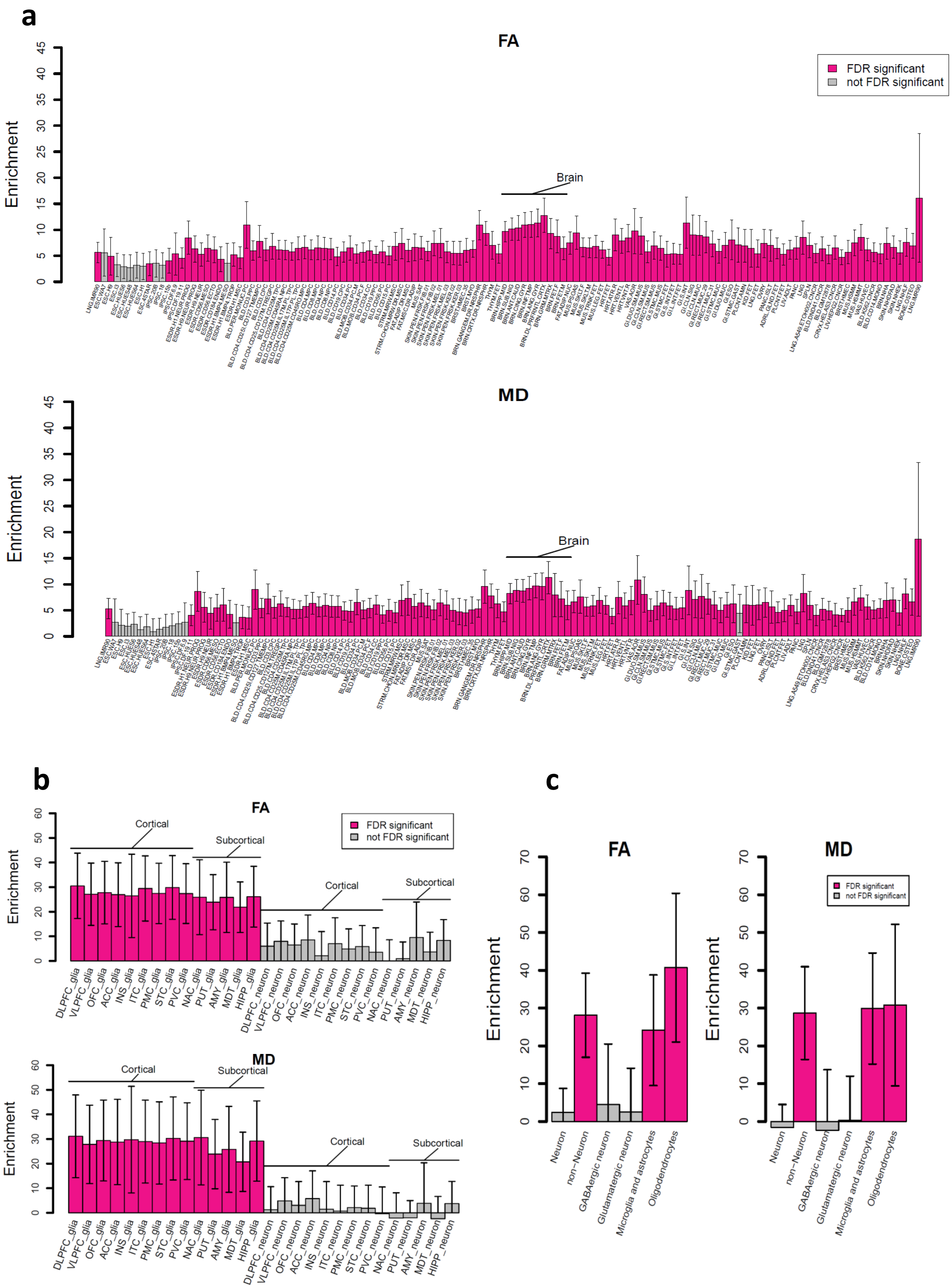


**Supplementary Figure 21:** (**A**) Heritability enrichment scores in regulatory elements of multiple tissue types; (**B**) Heritability enrichment scores in regulatory elements of cell types in multiple brain regions; and (**C**) Heritability enrichment scores in cell-type specific regulatory elements

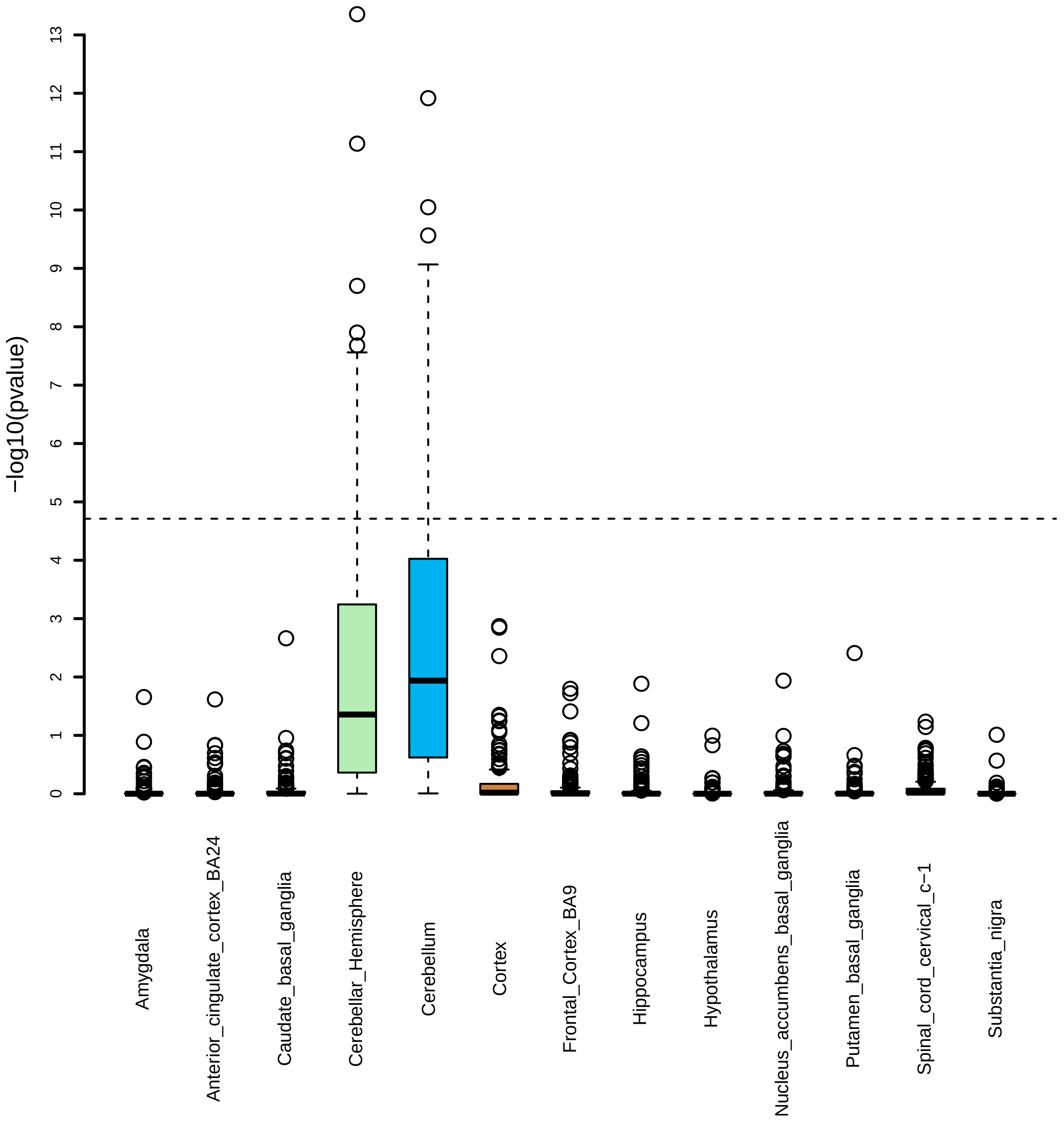


**Supplementary Figure 22:** MAGMA gene property analysis for UKB British discovery GWAS results (n=33,292 subjects) and 13 brain tissues. The 13 brain tissues were from GTEx v8 RNA-seq database. The tests above the horizontal dashed line are significant after Bonferroni correction.

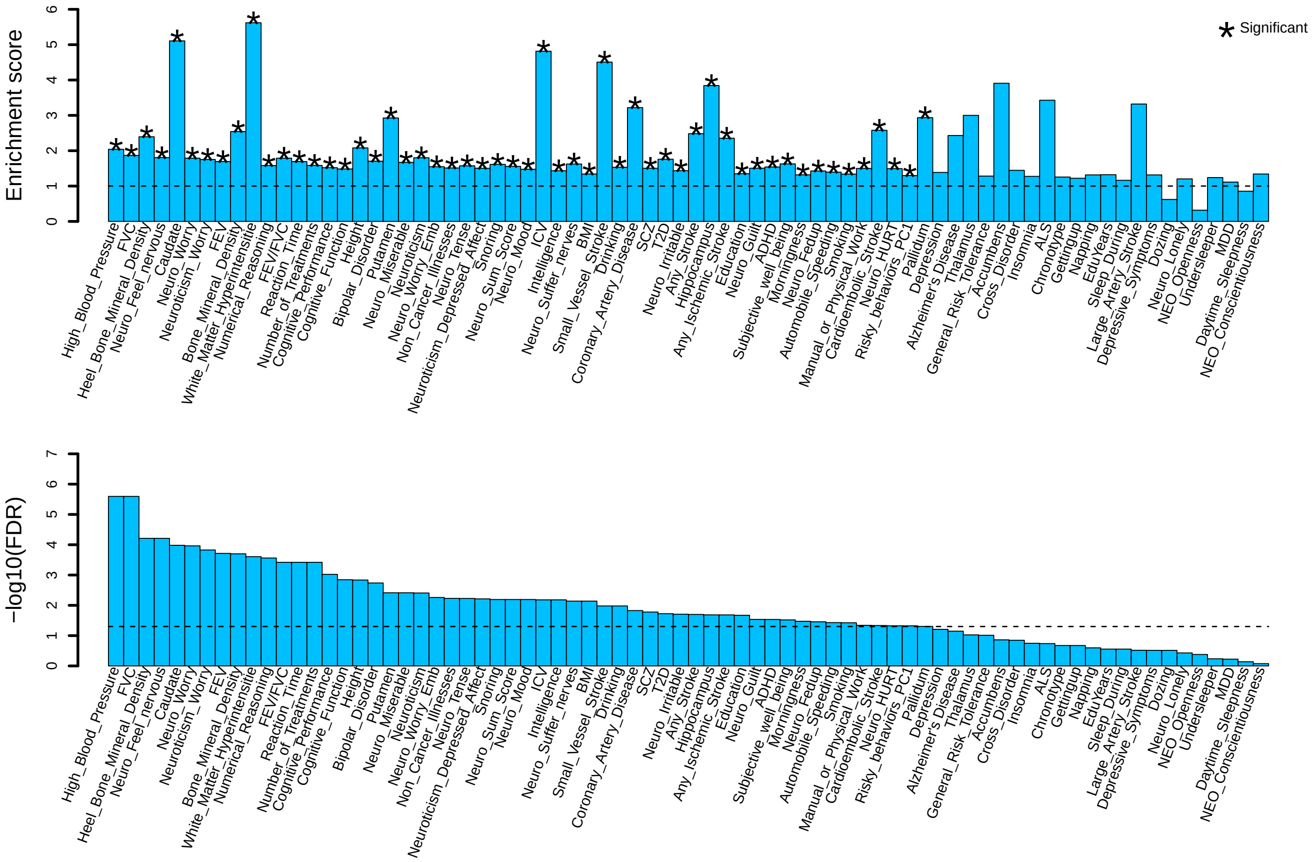


**Supplementary Figure 23:** Heritability enrichment scores and p-values in LDSC partitioned heritability analysis (with 95% confidence interval) for 76 complex traits. We treated DTI associated genes (n=33,292 subjects) as annotation and performed partitioned heritability enrichment (i.e., proportion of explained heritability/proportion of variants) analysis for other complex traits. The traits are ordered by their p-values.

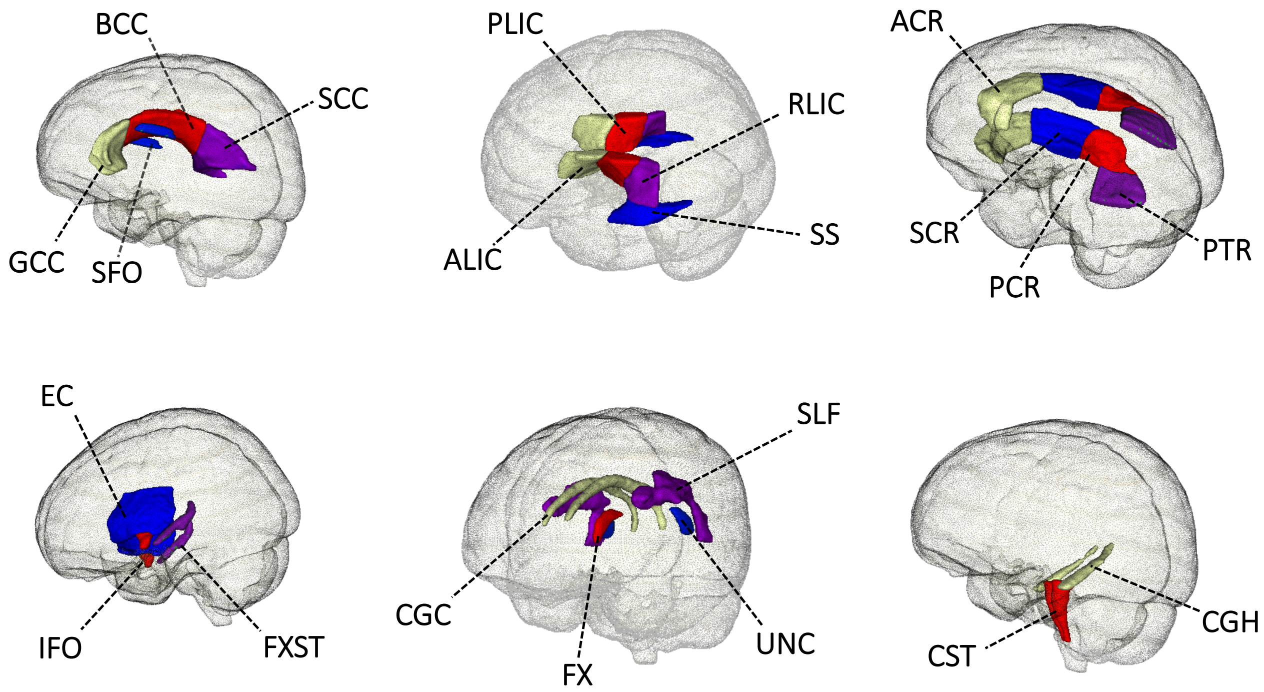


**Supplementary Figure 24:** Annotation of the 21 white matter tracts in human brain.

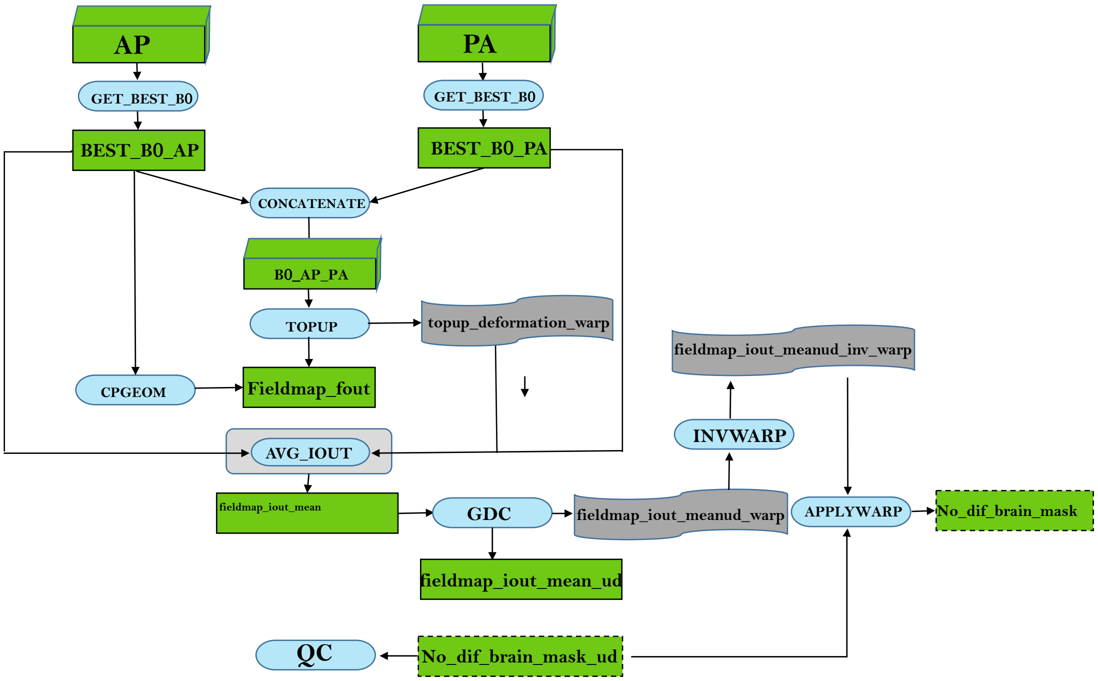


**Supplementary Figure 25:** Overview of fieldmap generation workflow used in this study to preprocess raw UK Biobank DICOM images.

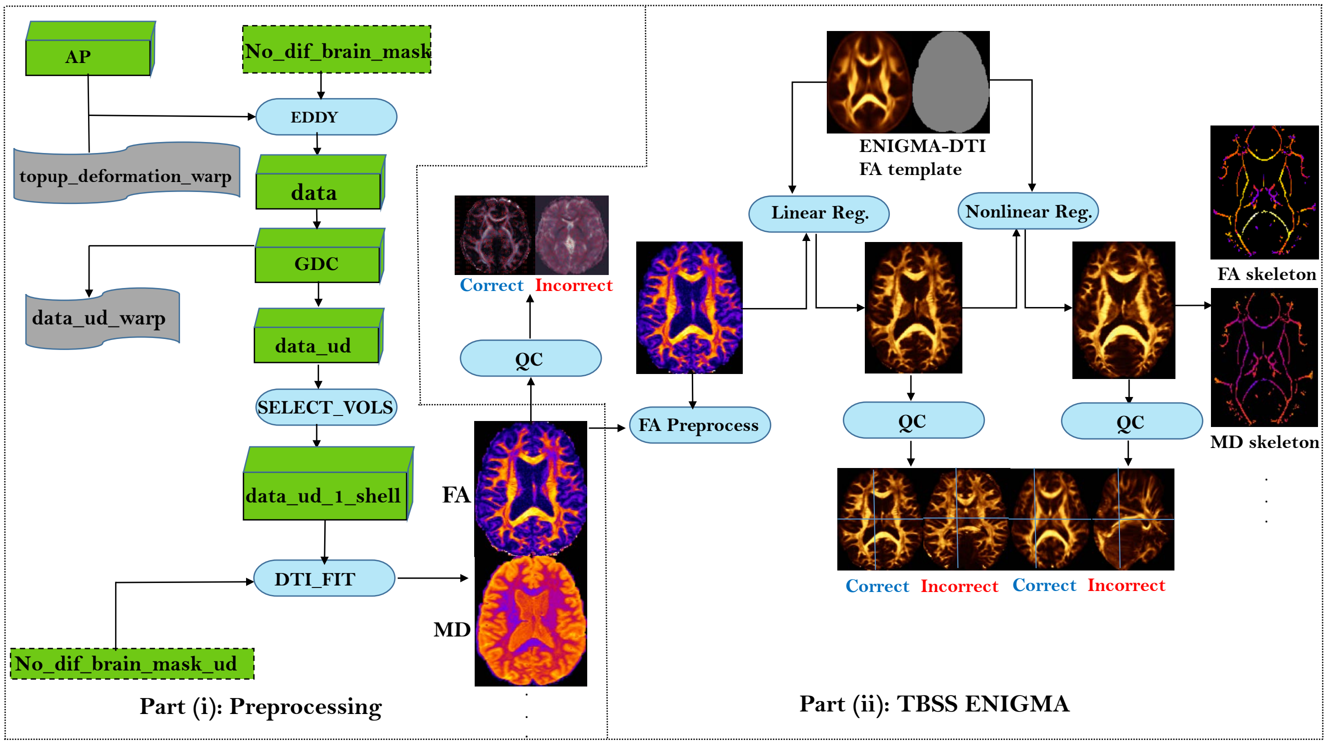


**Supplementary Figure 26:** Overview of the Eddy correction (left) and ENIGMA-DTI pipeline (right) used in this study.

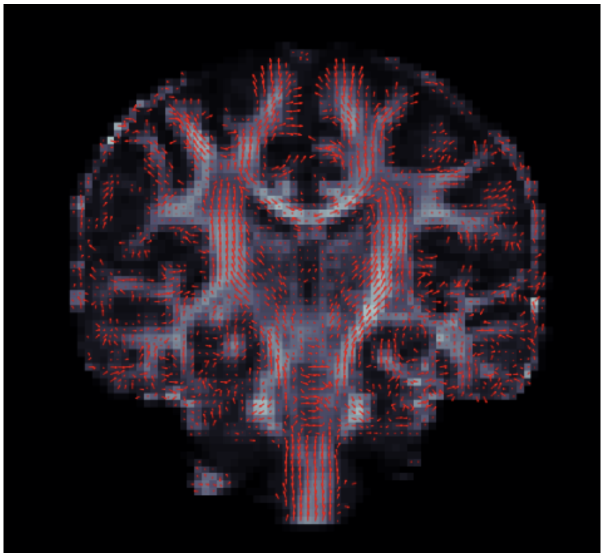

**Supplementary Figure 27**

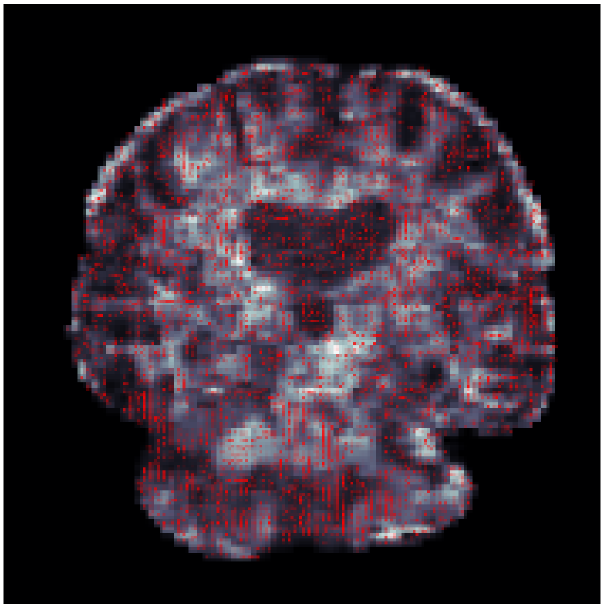

**Supplementary Figure 28**

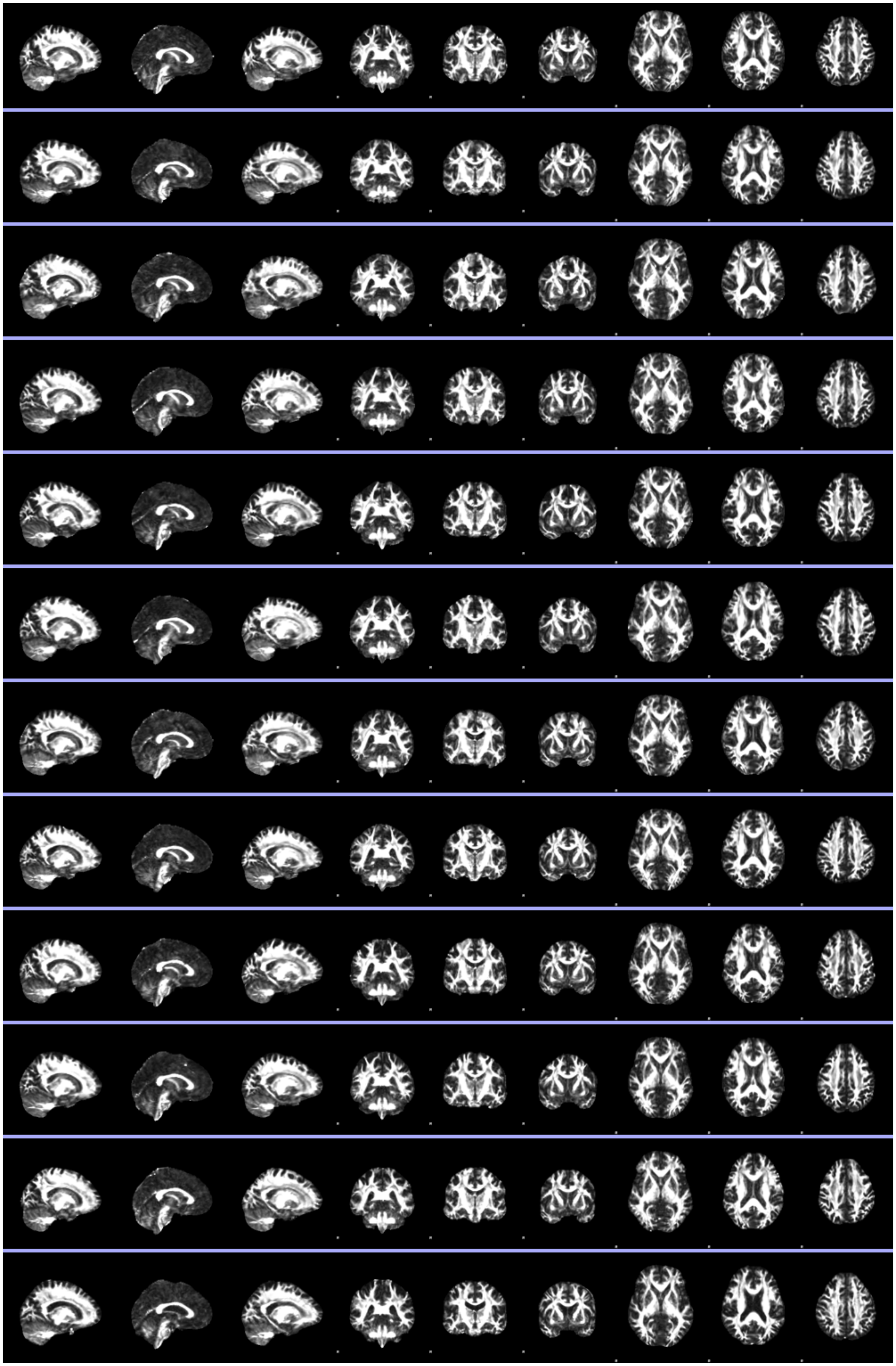

**Supplementary Figure 29**

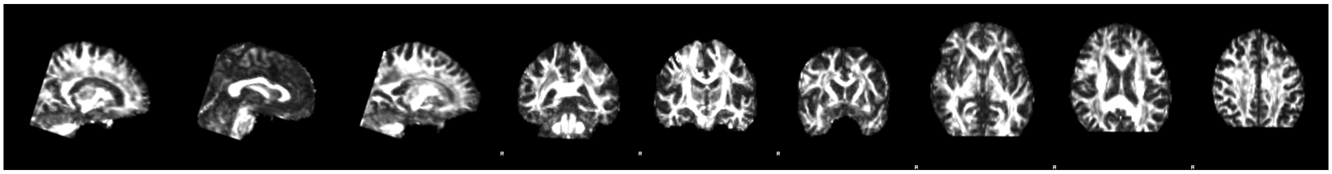

**Supplementary Figure 30**
